# Patient-derived liver biopsy organoids enable precision alcohol-associated liver disease modeling

**DOI:** 10.1101/2025.03.22.644563

**Authors:** Silvia Ariño, Laura Zanatto, Raquel A. Martínez-García de la Torre, Raquel Ferrer-Lorente, Jordi Gratacós-Ginès, Ana Belén Rubio, Martina Pérez, Beatriz Aguilar-Bravo, Guillermo Serrano, Stephen Atkinson, Zhengqing Xu, Paula Cantallops-Vilà, Laura Sererols-Viñas, Paloma Ruiz-Blázquez, Aina Rill, Juan José Lozano, Mar Coll, Idoia Ochoa, Silvia Affo, Anna Moles, Elisabetta Mereu, Ramón Bataller, Elisa Pose, Pau Sancho-Bru

**Affiliations:** Institut d’Investigacions Biomèdiques August Pi i Sunyer (IDIBAPS), Barcelona, Spain; Centro de Investigación Biomédica en Red de Enfermedades Hepáticas y Digestivas (CIBERehd), Barcelona, Spain; Liver Unit, Hospital Clinic, Barcelona, Spain; Computational Biology Program, CIMA University of Navarra, IdiSNA, Pamplona, Spain; Department of Metabolism, Digestion and Reproduction, Imperial College London; Department of Experimental Pathology, Institute of Biomedical Research of Barcelona, Spanish National Research Council (CSIC), Barcelona, Spain; Josep Carreras Leukaemia Research Institute, Barcelona, Spain; University of Barcelona (UB), Barcelona, Spain; Department of Electrical Engineering (TECNUN), University of Navarra, San Sebastian, Spain; Data Science and Artificial Intelligence Institute (DATAI), Universidad de Navarra, Pamplona, Spain

**Keywords:** organoid, liver, ALD, alcohol-associated liver disease, epithelial cells, precision medicine, steatotic liver disease, chronic liver disease, ductular reaction, cell plasticity

## Abstract

**Background & Aims:** Alcohol-associated liver disease (ALD) is a major cause of liver disease worldwide with scarce therapeutic options. Animal models poorly recapitulate advanced ALD precluding the development of new treatments. Organoids have emerged as a powerful human-based preclinical tool. However, current patient-derived liver organoids fail to recapitulate the epithelial heterogeneity and its generation requires liver surgical resections, thus limiting personalized disease modeling. Here, we report the development of organoids from liver needle biopsies (b-Orgs) from patients with ALD.

**Methods:** b-Orgs were generated from tru-cut biopsies from patients at early (n=28) and advanced (n=34) stages of ALD. b-Orgs were characterized by immunofluorescence, bulk and single cell RNA-sequencing and compared to parental tissues. b-Orgs were used to model ALD progression, identify pathogenic drivers, induce alcohol-associated hepatitis (AH) and evaluate response to prednisolone.

**Results:** Phenotypic and functional analysis of b-Orgs showed hepatocyte-enriched features. Single-cell RNA-sequencing revealed a heterogeneous cell composition comprising hepatocyte, biliary and progenitor populations, mirroring the epithelial landscape found in patients with advanced ALD. Moreover, b-Orgs preserved disease-stage features and allowed to identify the association of ELF3 with cell plasticity and disease progression. Finally, stimulation of b-Orgs with drivers of ALD induced pathophysiological features of alcohol-associated hepatitis, including ROS production, lipid accumulation, inflammation and decreased cell proliferation, which were mitigated in response to prednisolone.

Conclusions

Overall, we provide a human-based model that recapitulates epithelial complexity and patient specific features, allowing to identify drivers of cell plasticity and expanding organoid-based liver disease modeling for personalized medicine.

Impact and implications

Here, we describe the generation of biopsy-derived organoids (b-Orgs) from patients with liver disease. b-Orgs reproduce the liver epithelial cell composition found in patients’ liver tissue and are efficiently generated from different stages of the disease, providing a platform for patient-tailored disease modeling and drug testing.

## INTRODUCTION

Alcohol-associated liver disease (ALD) is a major cause of disability and premature mortality worldwide[1]. ALD comprises a broad spectrum; from steatosis to progressing inflammation and fibrosis to alcohol-associated hepatitis (AH), the most severe form of the disease. In addition to excessive alcohol consumption, factors such as genetic polymorphisms, gender or comorbidities participate in ALD pathogenesis and severity[2,3]. Therefore, individual-specific determinants are key elements involved in the progression of the disease. Besides abstinence and liver transplantation, available treatments are limited and focus on the management of alcohol intake and ALD complications. Indeed, AH still has a mortality rate of 20–50 % within the first three months. Despite several clinical trials on possible therapies, at present the only treatment available for these patients is corticosteroids, which unfortunately is only effective within the first months of treatment. Further, not all patients respond to them[4,5]. Thus, there is a clear need for a deeper understanding of the underlying disease mechanisms. However, the study of ALD and the development of new therapeutic options is limited by the lack of good preclinical models, as animal models poorly recapitulate human physiopathology and patients’ characteristics. Taken all together, these factors underscore the need to develop patient-derived *in vitro* systems to better recapitulate the disease condition and patient specificity.

Organoids are emerging as new tools to better understand and model molecular and cellular bases of disease[6]. Liver organoids have shown a significant potential for translational research of liver disorders. In this regard, the biomedical potential of liver organoids has been proven mainly with pluripotent stem cell-derived organoids, which are useful to model steatotic liver disease, drug toxicity and viral infection[7–11]. Patient-derived liver organoids have been successfully generated from diverse genetic liver disease conditions and cancer, recapitulating pathophysiological processes, and retaining patient-specific genetic backgrounds[12,13].

Organoids derived from liver cancer such as hepatocellular carcinoma and cholangiocarcinoma preserve the disease-related cancer mutations and drug response[14–18]. In addition, organoids generated from inherited genetic disorders such as Alagille syndrome[19] or alpha-1-antitrypsin deficiency[19,20] recapitulate patient-specific pathogenic dysfunction. However, few efforts have been made to develop organoids from acquired chronic liver diseases[21,22].

Patient-derived organoid generation mostly relies on the dissociation of resected liver tissue samples harvested at advanced stages[12,13], thus limiting the attainment of organoids from earlier stages of the disease. In addition, most studies generate cholangiocyte organoids, which present a homogeneous culture of biliary cells and do not recapitulate the different epithelial populations of the liver, as they lack the hepatocyte fraction[6,11,19–21,23]. In this regard, the establishment of human-derived organoids with hepatocyte features has only been accomplished using isolated fetal and non-disease primary human hepatocytes[24–26]. Only hepatocellular carcinoma tumoroids established from liver resections have proven capable of expressing robust phenotypic and functional hepatocyte traits[17]. Nonetheless, the generation of human organoids from chronic liver disease patients containing a hepatocyte-like population remains to be tested.

In this study we describe the generation of patient-derived liver organoids from intact tru-cut needle biopsies (b-Orgs). b-Orgs recapitulate the liver epithelial heterogeneity, preserve patient’s disease-stage features and are useful for patient-specific AH modeling.

## METHODS

### Data availability

Bulk and single cell RNA-sequencing datasets generated as part of this study are at the Gene Expression Omnibus and are publicly available as of the date of this publication. Accession number for bulk-RNA sequencing data is GSE272014. Accession number for scRNA-seq data is GSE273146.

### Human liver samples

Tru-cut liver needle-biopsies from early (n= 28) and advanced (n= 34) stages of ALD were used for organoid generation. Patients were considered at early stage when presenting compensated cirrhosis and fibrosis in pre-cirrhotic stages, whereas patients with decompensated cirrhosis or AH were stratified as advanced stage. The clinical and biochemical parameters of these patients are shown in Supporting Table S1.

Standard liver organoids were generated from non-tumour liver resections obtained at the time of transplantation from patients with advanced ALD.

All liver biopsies and resections used for organoid generation were selected from patients admitted to the Liver Unit of the Hospital Clinic of Barcelona. Informed consent was obtained from all the patients and the study was approved by the Ethics Committee of the hospital.

### Statistical Analysis

Experimental data are presented as mean values ± SEM. All parameters analyzed with parametric tests followed normal distribution by Anderson-Darling, D’Agostino & Pearson, Shapiro-Wilk and/or Kolmogorov-Smirnov tests. Non-normal distributed conditions were analysed by non-parametric tests as indicated in the figure legend. For normal distributed data, paired or unpaired two-tailed t-test were used when 2 groups were compared. For non-normal distributed data, Mann-Whitney test (non-paired samples) or Wilcoxon matched-pairs signed rank (paired samples) were used when 2 groups were compared; Kruskal-Wallis (non-paired samples) and Friedman (paired) tests with Dunn’s post-test were used to compare more than two groups. Two-way ANOVA with Sidak’s multiple comparisons was used for multiple groups comparison. Correlation coefficients (R) and p-values (*p*) were calculated by Pearson correlation test and linear functions were inferred by linear regression. Statistical analyses were performed using GraphPad v8.0.1 software.

All the other material and methods are described in the Supplementary Data.

## RESULTS

### Derivation of liver organoids from needle-biopsies at different stages of ALD

To generate organoids from different stages of the disease, liver biopsies were obtained from patients with ALD (28 from early and 34 from advanced stages). Patients presenting compensated cirrhosis and fibrosis in pre-cirrhotic stages were defined as early stage, whereas patients with decompensated cirrhosis or AH were stratified as advanced stage (Table S1). Needle biopsy fragments of 2-4mm were embedded in Matrigel and cultured with medium conditions for human hepatocyte[26] (HEP) and biliary[19,27] (BEC) organoid generation (Figure 1A). As early as two days after biopsy embedding, b-Orgs from both culture conditions started to bud (Figure 1B and 1C). b-Orgs were mechanically dissociated from the biopsy and subsequently plated in Matrigel for expansion (Figure 1C).

**Figure 1.**
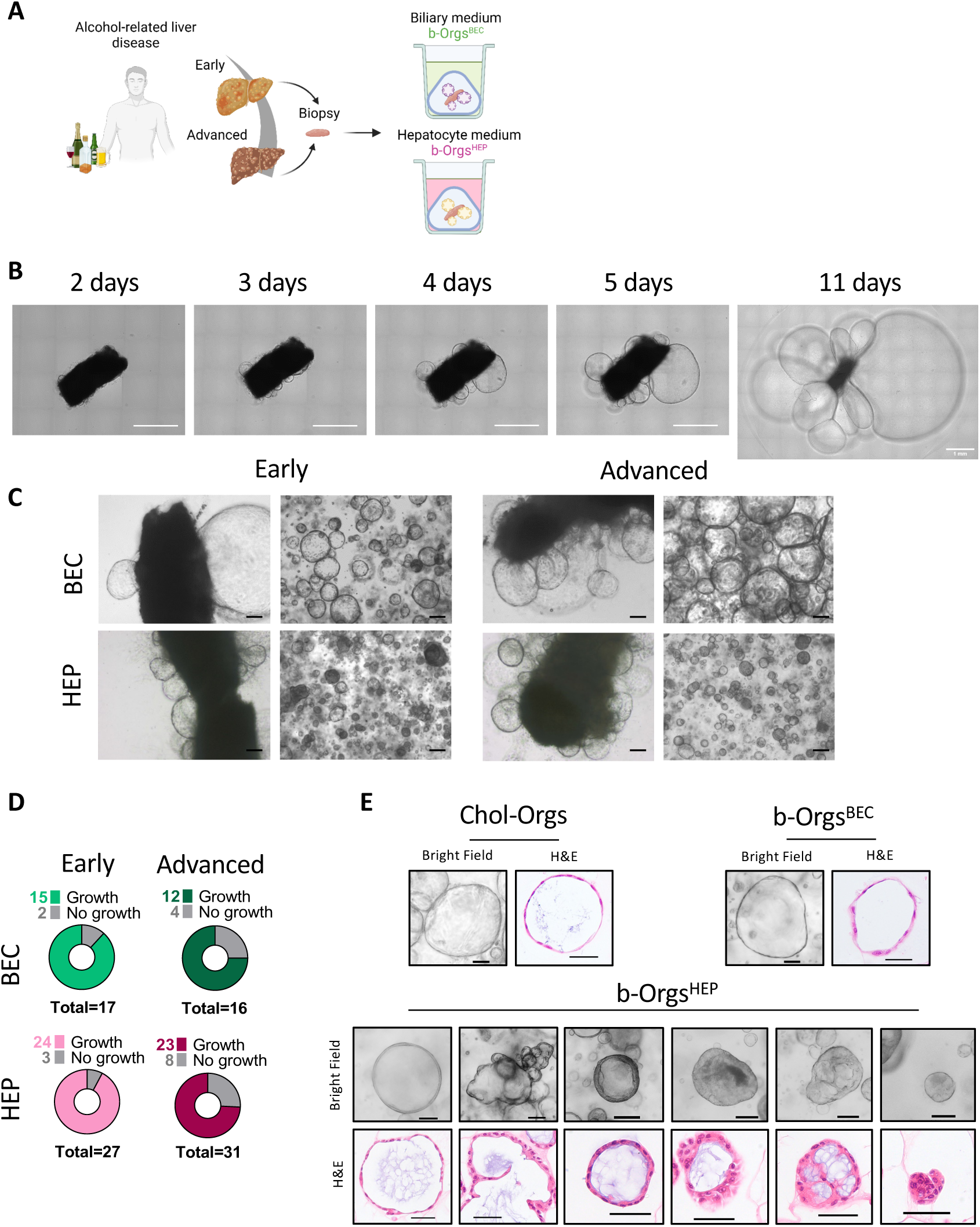
Derivation of liver organoids from needle-biopsies at different stages of ALD. **(A)** Scheme of the experimental procedure for organoid generation from needle biopsies. **(B)** Representative serial images of biopsy culture with budding organoids at the indicated time points. Scale bar = 1mm. **(C)** Representative images of biopsies and dissociated organoids from early and advanced ALD stages using HEP and BEC culture conditions. Scale bar = 200µm. **(D)** Efficiency of organoid generation considering disease stage and culture conditions (n=17 BEC early; n=16 BEC advanced; n=27 HEP early; n=31 HEP advanced). **(E)** H&E and bright field representative images of the different morphologies found in Chol- Orgs and b-Orgs. Scale bars = 100 µm (bright field) and 50 µm (H&E). ALD, alcohol-associated liver disease; b-Orgs, biopsy-derived organoids; Chol-Orgs, cholangiocyte organoids.

The success rate of b-Org generation was higher in patients with early ALD, with an efficiency of 88% with BEC medium and 89% using HEP medium versus 75% and 74% for BEC and HEP culture conditions respectively in advanced ALD (Figure 1D). While b-Orgs in BEC medium (b-Orgs^BEC^) grew as round cystic monolayers resembling the widely reported cholangiocyte organoids (Chol-Orgs) obtained by the dissociation of liver resections, b-Orgs generated with HEP medium (b-Orgs^HEP^) displayed heterogeneous morphologies, ranging from cystic to compact structures with spherical and multilobulated morphologies (Figure 1C and 1E and Supplementary Figures 1A, 1B and 1C). Moreover, b-Orgs^HEP^ showed a diversity of morphological structures comprising organoids with thick epithelial layers, multiple cavities or “cup-shaped” morphology (Figure 1E and Supplementary Figure 1A). Organoid morphology was patient-dependent (Supplementary Figure 1B and 1C), with no correlation between morphology and disease stage, thus suggesting that individual characteristics of the patient or tissue of origin may influence b-Org morphology.

Altogether, these results indicate that organoids can be successfully generated using a limited amount of liver tissue with minimal manipulation, enabling the *in vitro* modeling of the whole spectrum of ALD.

### Biopsies can be maintained long-term in culture while generating organoids

After disaggregating the budding organoids for subsequent expansion, biopsies were re-plated in Matrigel. We observed that a new generation of organoids budded from the re-plated biopsies after 2-4 days (Figures 2A and 2B). Successful production of new generations of b-Orgs could be achieved every 20 days for up to nine re-plating steps (Figure 2C, 2D and Supplementary Figure 2A and 2B). Although the production of b-Org generations was slower at advanced re-plating steps, biopsies could be maintained in culture for up to five months (Figure 2D). In addition, biopsies could be cryopreserved at any time for future b- Org generation (Supplementary Figure 2C). In parallel, b-Orgs could be expanded or cryopreserved. b-Orgs did not show morphological alterations among successive generations and passages (Figure 2B and Supplementary Figures 2B and 2D). Likewise, gene expression levels of biliary (*KRT7* and *EPCAM)* and hepatocyte (*ALB* and *CYP3A4)* markers were maintained across b- Org generations cultured in both BEC and HEP conditions (Figure 2E). When considering individual b-Org lines, we did not observe significant transcriptomic changes in the expression of liver cell markers between first, second and third generations of b-Orgs^HEP^ (Figure 2F). Moreover, there were no differences in liver cell markers between passages of b-Orgs^HEP^ (Figures 2G). We performed a weighted correlation network analysis (WGCNA) of b-Org^HEP^ transcriptomic data to characterize changes in functionality across generations and passages. Only a total of 174 genes distributed in three modules were significantly changed over time (Supplementary Figure 3E), accounting for a few biological processes which were mainly involved in ion homeostasis and morphogenesis (Supplementary Figure 3F).

**Figure 2.**
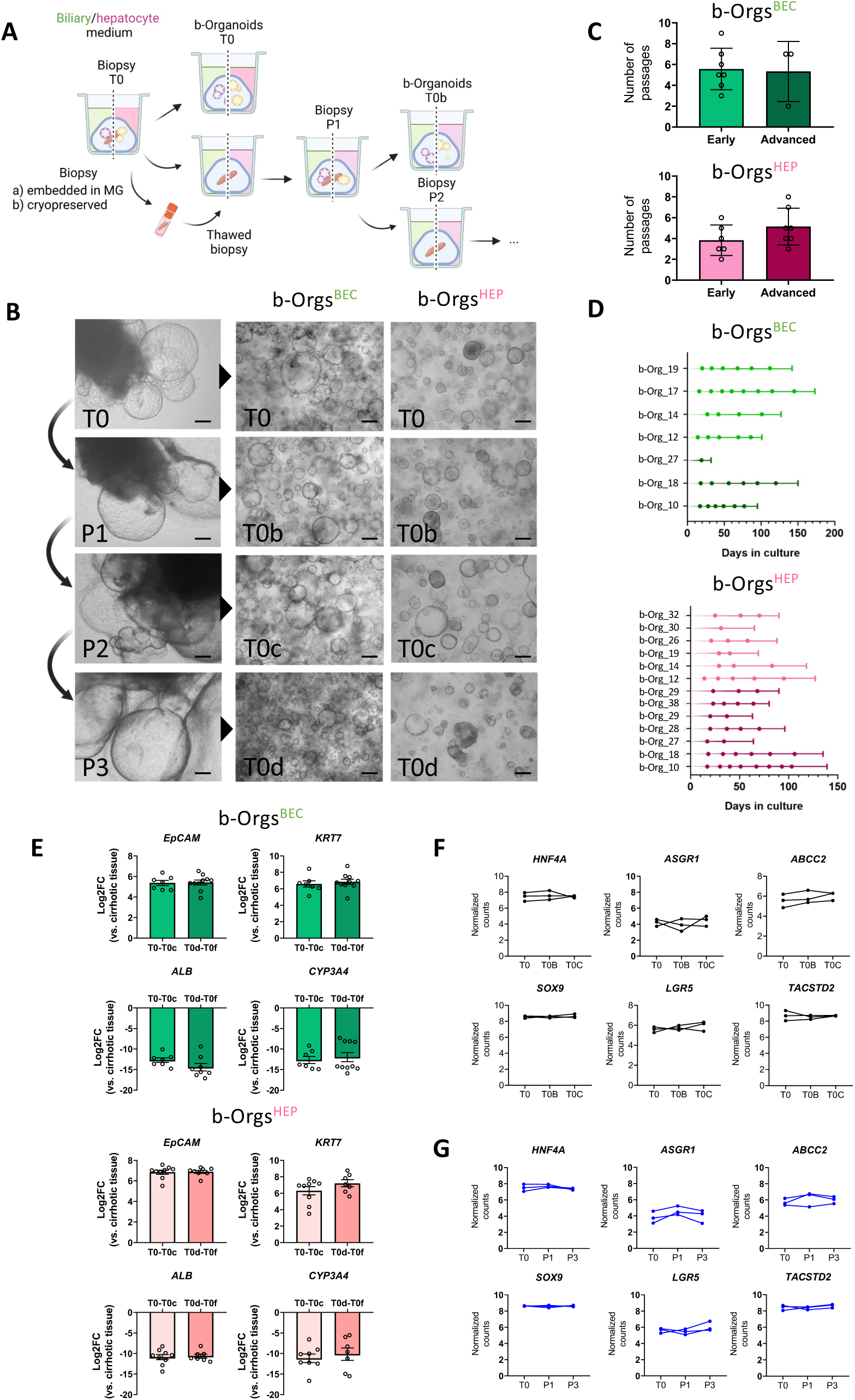
Biopsies can be maintained long-term in culture while generating organoids. **(A)** Scheme of the experimental procedure for biopsy maintenance in culture and b-Org generation. **(B)** Number of times that biopsies can be replated while generating b-Orgs (n=7, BEC; n=6-7 HEP). **(C)** Representative images of replated biopsies with budding organoids and successive b-Org generations. Scale bar = 200 μm. **(D)** Summary of the timeline of biopsies that were maintained in culture until no more organoids were generated. Each dot represents the time point when biopsies were replated, and budding organoids were separated and grown independently from biopsies. **(E)** Gene expression levels of biliary/progenitor (*EpCAM*, *KRT7*) and hepatocyte (*ALB*, *CYP3A4*) markers comparing early (T0-T0c) and late (T0d-T0f) generations of b-Orgs. At least, n=3 per group. **(F)** Normalized count levels for paired successive b-Org^HEP^ generations (n=3). **(G)** Normalized count levels for paired successive b-Org^HEP^ passages (n=3). All data are presented as mean ± SEM, no significance was determined by One-way ANOVA with Tukey’s multiple comparison (*ASGR1, SOX9, LGR5* and *TACSTD2* in F and G, and *ABCC2* in G), One-way ANOVA with Friedman test (*ABCC2* in F) and Two-way ANOVA with Sidak’s multiple comparison (E). b-Orgs, biopsy-derived organoids.

These findings show that needle-biopsies can be maintained under defined culture conditions and can give rise to patient-derived organoids that preserve stable transcriptomic and morphological characteristics over time.

### b-Orgs present enriched liver-like and hepatobiliary features

To date, reported patient-derived organoids have been obtained by tissue dissociation, and their cellular content is limited to a biliary phenotype, therefore being consensually designated as cholangiocyte organoids (Chol-Orgs)[6]. Given that Chol-Orgs are the existing model of patient-derived organoids from non-tumoral liver origin, we generated Chol-Orgs by tissue dissociation from ALD liver resections[22], and used them as a reference for b-Org characterization.

First, we performed bulk RNA-sequencing of b-Orgs and Chol-Orgs. Importantly, both b-Orgs^HEP^ and b-Orgs^BEC^ showed transcriptomic differences when compared to Chol-Orgs, as observed by principal component analysis. Notably, while b-Orgs^HEP^ showed the highest transcriptomic changes when compared to Chol-Orgs, b-Orgs^BEC^ also presented important differences with respect to Chol-Orgs, even when cultured with the same medium composition. These results indicate that the procedure for organoid generation has a direct impact on organoid phenotype, which is accentuated by the composition of the medium (Figure 3A).

**Figure 3.**
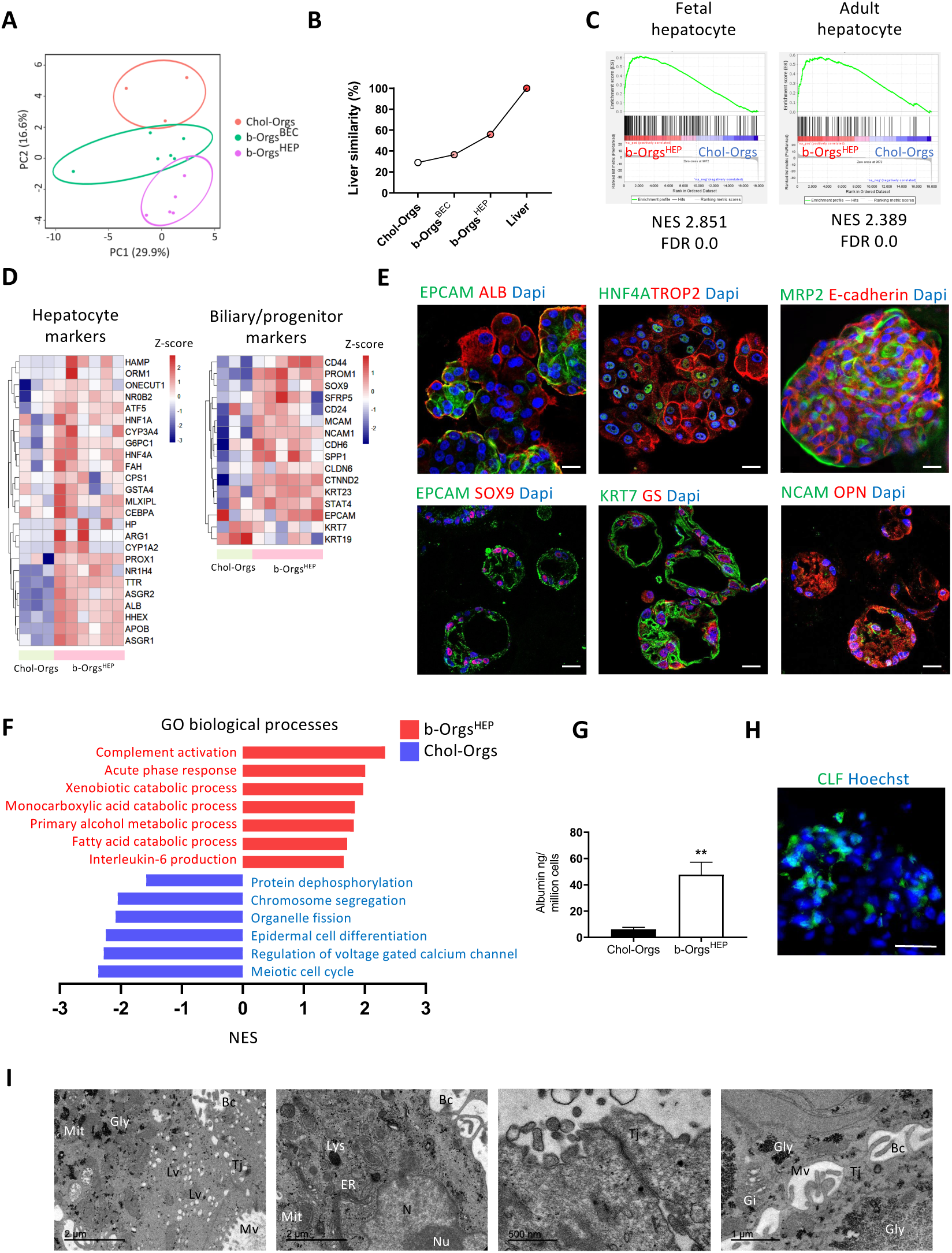
b-Orgs present enriched liver-like and hepatobiliary features. **(A)** PCA depicting transcriptomic differences among organoid groups. **(B)** Percentage (%) of liver similarity of cholangiocyte organoids and b-Orgs calculated using LiGEP algorithm[28,29]. **(C)** GSEA of hepatocyte signatures in b-Orgs^HEP^ versus Chol-Orgs using the pre-ranked mode. **(D)** Heatmaps depicting scaled expression levels of hepatocyte and biliary/progenitor genes in Chol-Orgs and b-Orgs^HEP^. **(E)** Representative images of b-Orgs^HEP^ stained for hepatocyte (ALB, HNF4A, MRP2, GS), progenitor/biliary (EPCAM, TROP2, SOX9, KRT7, NCAM, OPN) and epithelial (E-cadherin) markers. Scale bar = 20 µm. **(F)** Significantly enriched GO biological processes in b-Orgs^HEP^ (red) and Chol-Orgs (blue) using GSEA pre-ranked mode. **(G)** Albumin levels measured in organoid supernatant after 72 hours and normalized to one million cells (n=3 Chol-Orgs, n=15 b-Orgs^HEP^). **(H)** Representative image of CLF incorporation by b-Orgs^HEP^. Scale bar = 50 µm. **(I)** TEM images of b-Orgs^HEP^ showing mitochondria (Mit), glycogen (Gly), lipid vacuoles (Lv), bile canaliculi (Bc), microvilli (Mv), tight junctions (Tj), lysosome (Lys), endoplasmic reticulum (ER), nuclei (N), nucleolus (Nu) and Golgi apparatus (Gi). Data is presented as mean ± SEM, \*\**p* < 0.01 as determined by Mann-Whitney test in h. b-Orgs, biopsy-derived organoids; Chol- Orgs, cholangiocyte organoids; CLF, Cholyl-Lys-Fluorescein; DEGs, differentially expressed genes; GSEA, gene set enrichment analysis; GO, Gene Ontoloty; GS, glutamine synthetase; NES, normalized enrichment score; OPN, osteopontin; TEM, transmission electron microscopy.

Next, we used the transcriptomic data to interrogate the liver-specific gene expression panel (LiGEP) algorithm[28,29], which evaluates the degree of differentiation of hepatocyte *in vitro* models and their similarities to liver tissue. While Chol-Orgs presented a liver similarity score of 29.03%, b-Orgs revealed similarities of 36.56% and 55.91% under BEC- and HEP-culture conditions, respectively (Figure 3B).

Phenotypically, b-Orgs^HEP^ and b-Orgs^BEC^ showed higher normalized enrichment score (NES) values for biliary and hepatocyte signatures[30–33] when compared to Chol-Orgs (Figure 3C and Supplementary Figure 3A). However, b-Orgs^HEP^ were enriched in hepatocyte signatures in comparison to b-Orgs^BEC^ (Supplementary Figure 3A). Thus, considering that b-Orgs^HEP^ showed the highest similarity score to liver tissue and increased hepatocyte-like features over b- Orgs^BEC^, we further characterize them.

As expected, b-Orgs^HEP^ displayed hepatocyte markers such as *ALB*, *APOB* or *TTR* among the most differentially expressed genes (DEGs) when compared to Chol-Orgs, whereas genes involved in tissue development and oncogenesis were the most downregulated (Supplementary Figure 3B).

Remarkably, b-Orgs^HEP^ showed increased expression of hepatocyte canonical markers (i.e., *HNF4A*, *ASGR1*, *NR1H4*, *PROX1)* but also of progenitor/biliary markers such as *PROM1*, *SOX9* or *KRT23* when compared to Chol-Orgs (Figure 3D). In agreement with transcriptomic data, immunofluorescence analysis of b- Orgs^HEP^ confirmed the presence of cells expressing albumin, glutamine synthetase and HNF4A but also cells expressing EpCAM, SOX9, KRT7, TROP2, NCAM and osteopontin (OPN) (Figure 3E). Functional analysis revealed the upregulation of pathways related to hepatocyte inflammation and metabolism, such as complement activation and acute phase response or xenobiotic and lipid catabolism (Figure 3F and Supplementary Figure 3C). In addition, b-Orgs^HEP^ produced albumin (Figure 3G), formed functional bile canaliculi, stained with MRP2, which incorporated the bile acid analogue cholyl-L-lysyl-fluorescein (CLF) (Figures 3E, 3H and 3I) and stored glycogen (Figure 3I and Supplementary Figure 3D). Moreover, lipid accumulation and hepatocyte structure were revealed in b-Orgs^HEP^ by transmission electron microscopy (Figure 3I).

Mesenchymal signatures[32] were also enhanced in b-Orgs^HEP^ compared to both b-Orgs^BEC^ and Chol-Orgs (Supplementary Figures 3A and 3E). We detected some epithelial cells expressing vimentin and a small population of PDGFRB+ EPCAM- mesenchymal cells within the culture of several organoid lines (Supplementary Figure 3F).

These results indicate that b-Orgs^HEP^ display higher similarity to the liver tissue and enhanced hepatocyte features than Chol-Orgs.

### b-Orgs^HEP^ recapitulate the liver epithelial heterogeneity

Next, we aimed to evaluate the cell composition of the b-Orgs^HEP^. As shown in Figure 4A, b-Orgs^HEP^ were formed by cells showing different morphologies and sizes. To get a deeper insight into cell heterogeneity, we performed single-cell RNA-sequencing (scRNA-seq) analysis of b-Orgs^HEP^ derived from four different patients at early (n=1) and advanced (n=3) stages (Supplementary Figure 4A). After quality control filtering, we obtained transcriptomic information for a total of 14,183 cells, with a median of 3,232 cells from each organoid line and a median of 4,921 genes detected per cell (Supplementary Figure 4B). After performing batch correction among samples and merging the analyzed organoids, we visualized eleven different clusters by uniform manifold approximation and projection (UMAP) clustering (Figure 4B). Clusters were manually annotated with their respective cellular identities using biological processes associated with cluster DEGs together with canonical markers.

**Figure 4.**
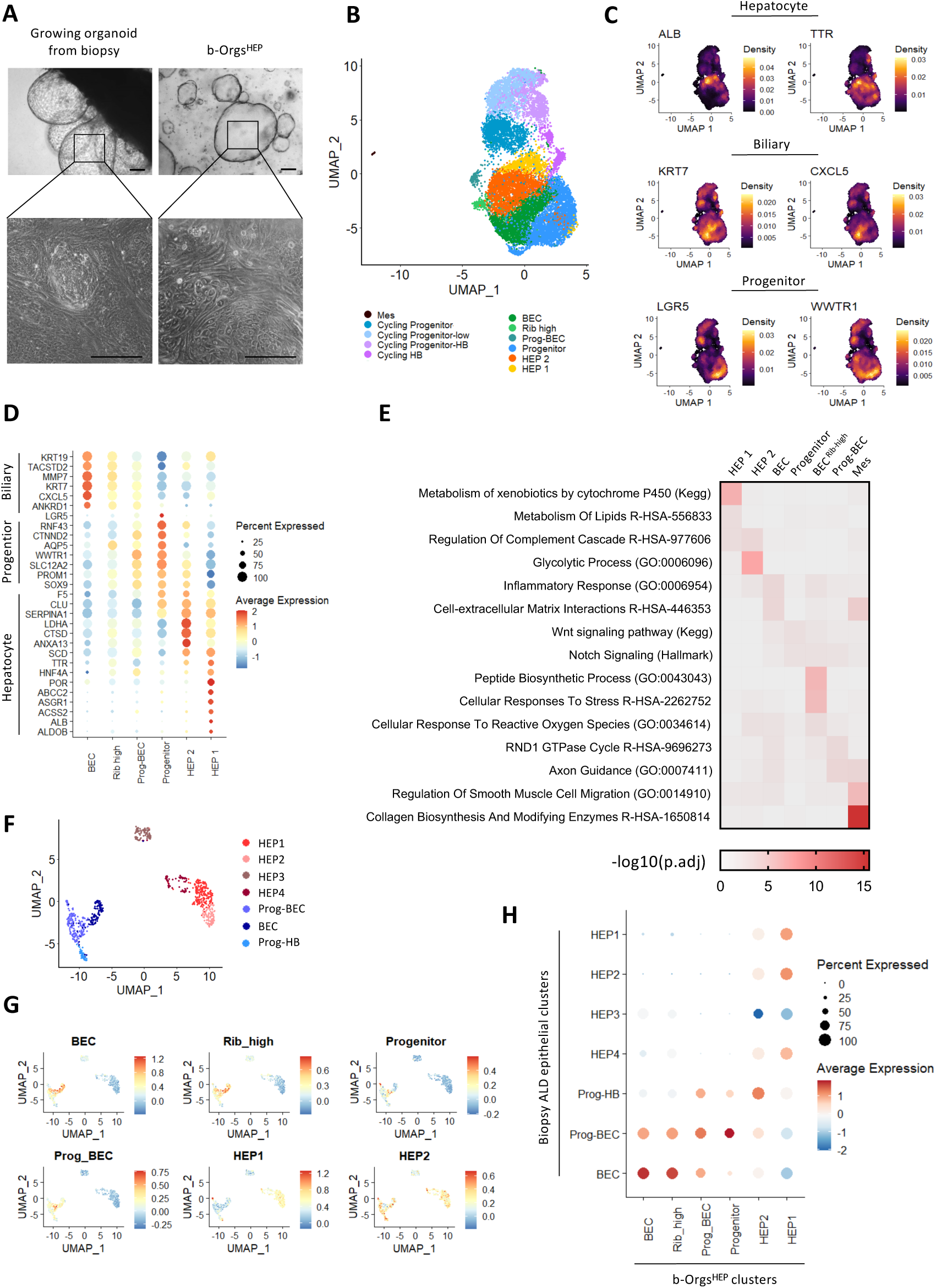
b-Orgs^HEP^ recapitulate the liver epithelial heterogeneity. **(A)** Representative bright field images of organoids when still budding from biopsies and after plating them separately. Scale bar = 100 µm. **(B)** UMAP visualization of the identified cell clusters forming b-Orgs^HEP^ (n=4). **(C)** Density expression of hepatocyte (*ALB*, *TTR*), biliary (*KRT7*, *CXCL5*) and stem (*LGR5*, *WWTR1*) markers visualized by UMAP plots. **(D)** Dot plot showing relative expression of marker genes for non-cycling hepatocyte, progenitor and biliary populations. **(E)** Heatmap displaying significantly enriched biological processes characterizing non-cycling clusters. **(F)** UMAP visualization of the epithelial compartment of biopsies from patients with ALD (n=2). **(G)** Enrichment of cluster- specific b-Org^HEP^ DEGs in epithelial clusters from ALD biopsies represented in UMAP as average expression in each cell. **(H)** Dot plot depicting enrichment (average expression) of cluster-specific b-Org^HEP^ DEGs in each epithelial cluster from ALD biopsies. Dot size represents the percentage of cells in every epithelial cluster of ALD biopsies enriched in b-Org^HEP^ signatures. ALD, alcohol-related liver disease; BEC, biliary; b-Orgs, biopsy-derived organoids; DEGs, differentially expressed genes; HB, hepatobiliary; HEP, hepatocyte; Prog, progenitor; Rib, ribosomal; UMAP, Uniform Manifold Approximation and Projection.

Although a small cluster of cells enriched in mesenchymal markers was detected (i.e., *PDGFRB*, *COL1A1, SPARC* or *IGFBP5*), most cells showed an epithelial phenotype expressing cell-cell interaction markers (i.e. *CDH1, OCLN or CLDN1*) (Supplementary Figure 4C).

We observed four clusters containing cycling cells with increased mitosis-related genes and G2/M and S phase scores, termed as cycling progenitor, cycling progenitor low, cycling progenitor-hepatobiliary (HB) and cycling HB (Supplementary Figures 4D, 4E and 4F). Pseudotime analysis revealed that cycling clusters of progenitor-like cells progressively evolved to an intermediate cycling HB population which eventually diversified into the main non-cycling clusters of cells with increased hepatocyte, progenitor and biliary characteristics (Supplementary Figures 4G and 4H).

Among non-cycling clusters, we found two clusters with hepatocyte-like features, HEP 1 and HEP 2 (Figure 4B). HEP 1 was highly enriched in mature hepatocyte genes such as *ALB* or *ABCC2,* metabolic processes such as xenobiotic detoxification (*ALDH3A1*, *ALDH2*, *CES2*) or lipid metabolism (*SCD*, *PPARG*, *ACADVL*) (Figures 4C, 4D and 4E). HEP 2 expressed *TTR*, *SERPINA1* and *ANXA13* but also progenitor genes (*SOX9, PROM1*) and hypoxia target genes (*NDRG1*, *BN1P3* or *VEGFA*) (Figures 4C and 4D). Functionally, HEP 2 was enriched in glucose metabolism (*LDHA*, *ALDOB*, *GAPDH*, *PKM*) and complement activation (*SERPING1*, *CFH*, *CFB*) (Figure 4E).

In addition, transcriptomic data revealed two clusters enriched in the progenitor markers *PROM1*, *SOX9*, *SLC12A2* and *WWTR1* (Progenitor and Progenitor- BEC) (Figures 4B, 4C and 4D). The Progenitor cluster was enriched in stem-related genes (*LGR5, RNF43*) and liver development and plasticity associated pathways including wnt and notch signaling. To the contrary, the Progenitor-BEC was enriched in biliary and progenitor markers, as well as axon guidance (*NTN4*, *EFNA5*) or RND1 GTPase signaling (*DEPDC2B*, *STMN2*) (Figures 4C, 4D and 4E). Therefore, the results suggest a transitioning phenotype between progenitor and hepatobiliary commitment.

Biliary genes such as *KRT7*, *KRT19* or *TACSTD2* were enriched in the BEC cluster. This cluster showed an enhanced pro-inflammatory phenotype (*CXCL5*, *CXCL6*, *IL18*) and an increased involvement in cell-extracellular matrix interactions (*ANXA1*, *ITGA3*, *CDH2*) (Figures 4C, 4D and 4E). In addition, we identified a small population with increased expression of cholangiocyte markers, but also enriched in ribosomal transcripts and stress-related processes (BEC^Rib-high^) (Figures 4B, 4D and 4E). Cells from all the clusters were present in all samples in variable proportions (Supplementary Figures 4I and 4J).

To evaluate to what extent the cellular heterogeneity of the organoids recapitulates the epithelial populations found in chronic liver diseases, we compared b-Org^HEP^ data with scRNA-seq data from the epithelial fraction of patients with advanced ALD. Interestingly, hepatocyte, progenitor and biliary clusters found in b-Orgs^HEP^ matched with the different epithelial clusters found in biopsies from patients with ALD (Figures 4F, 4G and 4H). These findings indicate that b-Orgs^HEP^ capture the epithelial landscape found in liver disease.

Altogether, these data reveal the epithelial composition of b-Orgs^HEP^, which recapitulates the epithelial heterogeneity found in the context of liver disease.

### b-Orgs^HEP^ recapitulate disease stage features

Given that needle-biopsies enable the generation of organoids from different stages of the disease, we aimed to explore whether b-Orgs^HEP^ preserve disease stage features observed in the parental tissue. To address this question, we performed bulk RNA-sequencing of b-Orgs^HEP^ and paired biopsies from patients at early and advanced stages of ALD and obtained the DEGs characterizing each disease stage. Gene Set Enrichment Analysis (GSEA) showed that DEGs from early-stage biopsies were significantly enriched in b-Orgs^HEP^ generated from these same patients. Likewise, b-Orgs^HEP^ from patients with advanced disease were enriched in DEGs from paired biopsies (Figure 5A). The core enrichment genes from this analysis were used to evaluate the biological functions and upstream regulators preserved by organoids at each disease stage.

**Figure 5.**
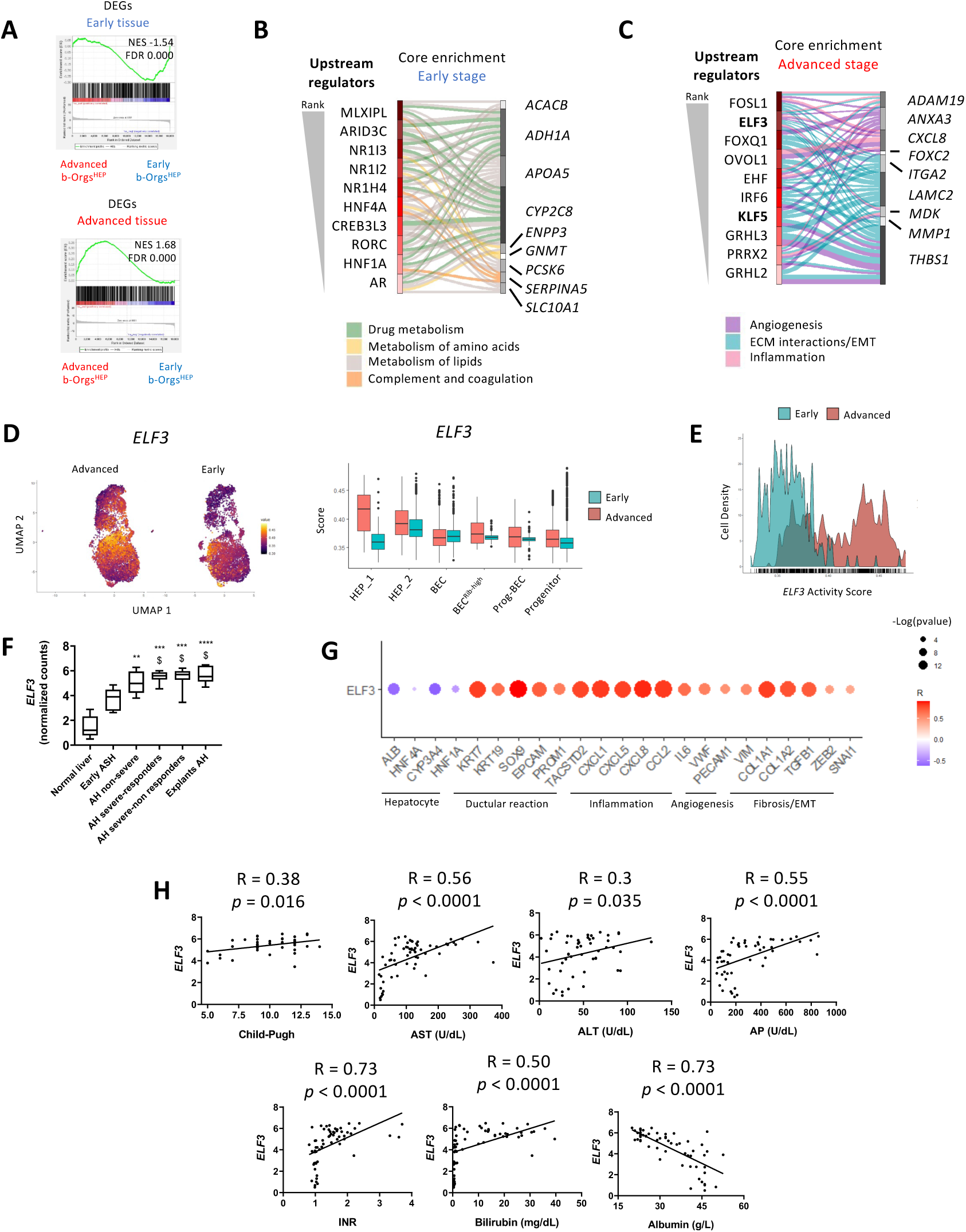
b-Orgs^HEP^ recapitulate disease stage features. **(A)** GSEA using DEGs from comparing biopsies at early (n=3) and advanced (n=3) ALD stages in pre-ranked b-Orgs^HEP^ from paired early-stage biopsies (n=3) versus b-Orgs^HEP^ from paired advanced-stage biopsies (n=3). **(B)** and **(C)** Alluvial plots showing top ten upstream regulators (left side of the plots, ranked from top to bottom by Chea3 tool) predicted to regulate genes belonging to the core enrichment gene set shared between b-Orgs^HEP^ and paired early-stage **(B)** and advanced-stage **(C)** biopsies. Representative genes from this core enrichment data set are shown on the right side of the plots. Upstream regulators and core enrichment genes are connected by biological processes in which the core enrichment genes are predicted to be involved. Each color represents a biological process. Some upstream regulators and core enrichment genes may be involved in more than one biological process. **(D)** On the left, UMAP visualization of predicted *ELF3* activity in early (n=1) and advanced (n=3) b-Org^HEP^ scRNA-seq data sets. On the right, box plots showing average activity score for *ELF3* in each of the non-cycling epithelial clusters, dividing samples in early and advanced ALD condition. Boxes represent interquartile range (IQR), and whiskers represent minimum and maximum values. Dots represent outliers. **(E)** Predicted *ELF3* regulon activity score distribution for each cell at early and advanced disease stages. **(F)** *ELF3* gene expression levels in total liver tissue of a cohort of patients with ALD. **(G)** Dot plot visualization of the correlation between the transcript levels of *ELF3* and hepatocyte, ductular reaction, inflammation, angiogenesis, fibrosis and EMT markers in a cohort of patients with ALD. The size of the dot represents the significance (p-value) and the color depicts the correlation coefficient (R) as determined by Pearson correlation test. **(H)** Correlations of *ELF3* mRNA expression in liver tissue from an ALD cohort with Child-Pugh score, international normalized ratio (INR) and with serum levels of alanine aminotransferase (ALT), aspartate aminotransferase (AST), alkaline phosphatase (AP), bilirubin and albumin. Correlation coefficients (R) and p-values (*p*) were calculated by Pearson correlation test and linear functions were inferred by linear regression. Data is presented as mean ± SEM; \*\**p* < 0.01, *** *p* < 0.001 and \*\*\*\**p* < 0.0001 versus Normal liver; $ *p* < 0.05 versus Early ASH, as determined by Kruskal-Wallis test with Dunn’s multiple comparison in F. AH, alcohol-related hepatitis; ALD, alcohol- associated liver disease; ASH, alcohol-associated steatohepatitis; b-Orgs, biopsy-derived organoids; DEGs, differentially expressed genes; EMT, epithelial to mesenchymal transition; GSEA, gene set enrichment analysis; NES, normalized enrichment score.

Core enrichment genes from early ALD stage were related to hepatocyte functions such as drug metabolism, amino acid and lipid metabolism, complement and coagulation (Figure 5B and Supplementary Figure 5A). In addition, among predicted upstream regulators we identified essential hepatocyte transcription factors involved in cell identity or functionality such as HNF4A or NR1I2 (Figure 5B). On the other hand, core enrichment genes in advanced stages of the disease were involved in injury-related processes such as inflammation, extracellular matrix remodeling and angiogenesis (Figure 5C and Supplementary Figure 5B), which were predicted to be driven by transcription factors related to cell damage, including ELF3, EHF and KLF5 (Figure 5C).

Next, we assessed whether b-Orgs could be used to identify molecular drivers of cell plasticity during disease progression. For this purpose, we applied the SimiC algorithm[34] to the b-Org^HEP^ scRNA-seq data set in order to infer the gene regulatory dynamics at the single-cell level between early and advanced disease conditions. SimiC indicated that the already-identified *ELF3* and *KLF5* had increased activity scores in the advanced ALD stage. Interestingly, whereas *KLF5* showed an increased activity in most clusters (Supplementary Figures 5C and 5D), *ELF3* activity was specifically enhanced in the HEP 1 cell population (Figure 5D). In addition, *ELF3* showed a bimodal activity score distribution with two cell populations displaying low and high *ELF3* activity (Figure 5E). These results suggest that *ELF3* could be regulating the hepatocellular compartment in ALD progression. To assess the clinical relevance of *ELF3*, we used whole liver transcriptomic data from a cohort of patients with ALD[35] to evaluate the correlation of *ELF3* expression with clinical parameters. We observed that *ELF3* expression increases with disease progression (Figure 5F) and correlates negatively with hepatocyte markers and positively with ductular reaction, inflammation, angiogenesis, fibrosis or epithelial-to-mesenchymal transition (EMT) (Figure 5G). Furthermore, as shown in Figure 5H, liver gene expression of *ELF3* positively correlated with Child-Pugh and analytical parameters of liver injury (i.e., AST, ALT, AP, serum bilirubin). Additionally, *ELF3* transcript levels showed a negative correlation with albumin concentration in serum and a positive correlation with the international normalized ratio (INR), denoting an association with impaired synthetic and coagulation function (Figure 5H).

Overall, these results show that b-Orgs recapitulate disease stage features at the basal level and indicate their potential for discovering potential disease drivers such as ELF3.

### b-Orgs model ALD progression and AH pathophysiology

ALD encompasses a broad spectrum of pathogenic features such as steatosis, inflammation and hepatocellular damage. Besides alcohol metabolites, other pathogenic drivers such as pro-inflammatory mediators and pathogen-associated molecular patterns (PAMPs) play a key role in the progression of ALD and, particularly, in the development of AH[4,36]. To mirror AH conditions, we treated b-Orgs^HEP^ for five days with a medium containing tumor necrosis factor-alpha (TNFa), interleukin 1-beta (IL1b), lipopolysaccharide (LPS) and ethanol (EtOH) (AH-medium). As shown in Figure 6A, b-Orgs^HEP^ from both early and advanced disease stages presented morphological changes after treatment with the AH- medium.

**Figure 6.**
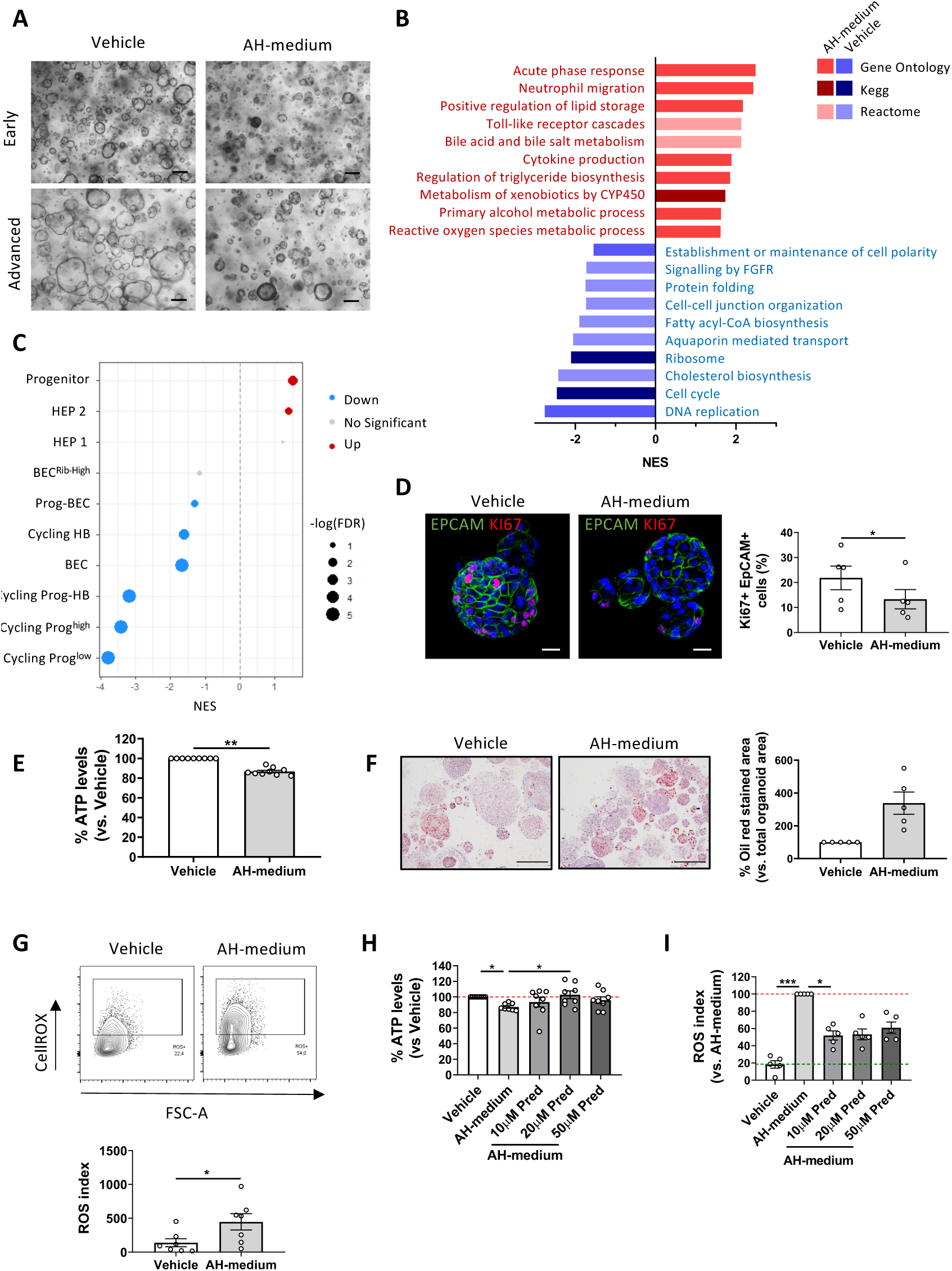
b-Orgs model ALD progression and AH pathophysiology. **(A)** Representative bright field images of early- and advanced-stage b-Orgs^HEP^ after a 5-day treatment with vehicle or AH-medium (10ng/mL IL1b, 20ng/mL TNFa, 100ng/mL LPS and 100mM EtOH). Scale bar = 200 μm. **(B)** Bar plot showing NES values for enriched biological processes in AH-medium treated b- Orgs^HEP^ (red bars) versus vehicle group (blue bars) (n=4). **(C)** Dot plot showing enrichment (NES values) of cluster-specific DEGs from b-Org^HEP^ scRNA-seq in b-Orgs treated with AH-medium and vehicle (n=4). Dots in red show positive enrichment in AH-medium treated group, dots in blue show negative enrichment and grey dots depict no significant changes. Dot size represents -log10(FDR) values. **(D)** Representative images of b-Orgs^HEP^ stained for EPCAM and the proliferation marker KI67 after treatment with vehicle and AH-medium. Average of KI67+ EPCAM+ cell quantification is shown as a percentage (%) relative to total nuclei (n=5). **(E)** ATP levels measured in vehicle and AH-medium treated b- Orgs^HEP^. ATP levels are presented as percentage (%) normalized to paired vehicle condition (n=9). **(F)** Representative images of b-Orgs^HEP^ stained for Oil red at day 5 after AH-medium treatment and vehicle condition. Quantification of Oil red organoid-stained area relative to total organoid area is shown as a percentage (%) (n=5). Scale bar = 200 μm. **(G)** Representative flow cytometry plots of ROS levels in treated and untreated b-Orgs^HEP^. Quantification of ROS index (ROS+ live cells multiplied by ROS median intensity of live cells) (n=7). **(H)** ATP levels measured in b-Orgs^HEP^ at day 5 of treatment with AH-medium and different concentrations of Pred added to the AH-medium during the last three days of treatment. Data are shown as percentage (%) versus paired vehicle (n=8). **(I)** ROS index determined in b-Orgs^HEP^ at day 5 of treatment with AH-medium and different concentrations of Pred added to the AH-medium during the last three days of treatment. Data are shown as a percentage (%) versus paired AH-medium treated samples (n=5). All data are presented as mean ± SEM; no significance or * *p*< 0.05, \*\**p* < 0.01, were determined by Wilcoxon matched-pairs signed rank test (E and F), Friedman test with Dunn’s multiple comparisons test (H and I), paired t-test (D and G). AH, alcohol-associated hepatitis; BEC, biliary epithelial cells; b-Orgs, biopsy-derived organoids; EtOH, ethanol; HB, hepatobiliary; HEP, hepatocyte; Pred, prednisolone; Prog, progenitor; ROS, reactive oxygen species.

Transcriptomically, organoids treated with the AH-medium upregulated pro-inflammatory processes such as acute phase response or cytokine production and pathways related to production of oxygen species, lipid accumulation and detoxification (Figure 6B). We confirmed the response to the AH medium by evaluating the gene expression of the senescence marker *CDKN1A*, pro-inflammatory mediators (*CCL2*, *CXCL8*) and markers of metabolic response to ethanol (*ADH1A*, *CYP2E1*, *PXR*) (Supplementary Figure 6A). On the contrary, treatment with AH-medium induced a downregulation of cell-cycle and homeostatic processes including DNA replication, cell-cell junction organization or protein folding (Figure 6B). Importantly, expression of genes involved in pro-inflammatory and oxidative stress response showed the same trend in treated organoids and AH tissue, thus reinforcing the idea that organoids recapitulate the liver tissue profile from patients with AH (Supplementary Figure 6B). Interestingly, b-Orgs^HEP^ treated with AH-medium overexpressed progenitor markers (*LGR5*, *PROM1*, *STAT4*) and were enriched in characteristic DEGs of the Progenitor-like cluster. In addition, treated b-Orgs^HEP^ clearly downregulated the expression of biliary markers (*EPCAM*, *KRT19*) and DEGs of BEC and cycling clusters identified in the organoid scRNA-seq data, thus suggesting an impact of AH- medium on organoid epithelial plasticity (Figure 6C and Supplementary Figure 6C).

We functionally confirmed the transcriptomic findings *in vitro*. The AH-medium induced a decrease in b-Org^HEP^ proliferation as indicated by a reduction of Ki67 staining (Figure 6D) and ATP levels (Figure 6E) with no changes in cell death (Supplementary Figure 6D). Besides, increased lipid accumulation and reactive oxygen species (ROS) were revealed by Oil Red staining and flow cytometry analysis, respectively, after treatment with AH-medium (Figures 6F and 6G). Notably, EtOH alone also reduced ATP levels while increasing ROS in treated b- Orgs^HEP^ (Supplementary Figures 6E and 6F).

Finally, we addressed whether b-Orgs^HEP^ could respond to prednisolone, the reference treatment for AH management. First, we treated b-Orgs^HEP^ at basal conditions with prednisolone, which induced a mild increase of ATP levels and a significant decrease of ROS (Supplementary Figures 6G and 6H). As shown in Figures 6H and 6I, prednisolone reversed the effects of AH-medium in b-Orgs^HEP^, increasing ATP and reducing ROS levels. Interestingly, different intensities of response to prednisolone were observed in each of the b-Org^HEP^ lines, denoting a potential patient-specific response (Supplementary Figure 6I).

These results indicate that b-Orgs^HEP^ respond to pathogenic drivers, recapitulating pathogenic processes involved in ALD progression and response to treatment.

## DISCUSSION

The development of advanced *in vitro* 3D systems recapitulating patients’ characteristics and disease-stage features is crucial to dissect disease pathophysiology and to identify new therapeutic targets[37]. Organoids have emerged as promising *in vitro* systems to model liver disease conditions. However, although continuous advances are being achieved in the field, patient-derived organoid generation is almost limited to end-stage liver resections and rendering organoids with a biliary phenotype, referred to as cholangiocyte organoids[12,13]. Here, we describe a novel approach enabling the generation of complex epithelial organoids derived from liver disease patients which we named b-Orgs. We show that b-Orgs recapitulate disease features and epithelial plasticity, identifying ELF3 as a potential driver of disease progression.

The methodology presented here enables the generation of organoids from tru-cut needle biopsies without enzymatic or mechanic disruption of the tissue. This is key for the organoid cellular composition and phenotype and for improving the success rates of organoid generation. This is particularly important when the source of starting material is limited, possibly helping to resolve the limited efficiency reported in previous reports[18] [16]. Moreover, a broader spectrum of disease stages and samples can be used with this procedure. Indeed, small intact fragments of end-stage liver resections can also give rise to b-Orgs with both HEP and BEC culture conditions, thus creating an *in vitro* platform that can cover the whole spectrum of the disease. We have thus set up a simple, fast, and reproducible procedure for organoid generation using a minimum amount of tissue sample.

*In vitro* maintenance and expansion of functional mature cells that faithfully resemble adult liver tissue has remained challenging, especially for hepatocytes.

Here we show that organoids directly generated from biopsies embedded in Matrigel present increased liver-like features with both BEC and HEP culture conditions when compared to Chol-Orgs. These results suggest that biopsies in culture may create a microenvironment that confers enhanced mature liver characteristics to organoids, which are maintained over time when b-Orgs are expanded. Functional enrichment analysis showed that liver-related processes such as detoxification, complement activation or lipid metabolism were increased in b-Orgs^HEP^—to the detriment of proliferation pathways—when compared to Chol-Orgs. Moreover, in contrast to the important expansion capacity of standard Chol-Orgs[19], b-Orgs’ successive passage is more limited, similarly to what has been previously reported for organoids from adult primary human hepatocytes[26]. These findings are consistent with previous studies stating that cell proliferation and maturation are negatively correlated and that hepatocytes can acquire hepatoblast features adapting their metabolic and proliferative status on demand with regenerative purposes[38,39]. In this regard, further studies should be performed to improve culture conditions, and overcome the limited proliferative capacity of b-Orgs while improving their maturation.

Up to now, patient-derived organoids described in the literature have been formed by biliary cells[40], thus limiting the study of the whole epithelial landscape found within the context of liver disease. Here we show by scRNA-seq that b-Orgs^HEP^ are integrated by several populations of epithelial cells expressing canonical markers of hepatocytes, cholangiocytes and/or progenitor cells. This is particularly important since ALD is characterized by plasticity events which determine the phenotype and function of hepatocyte and biliary populations. Indeed, hepatocyte loss of identity and reactivation of developmental pathways have been described as key pathogenic mechanisms involved in advanced ALD and poor patient outcomes[35,41–43], features that can be mimicked by b- Orgs^HEP^.

b-Orgs^HEP^ show two cell populations with hepatocyte traits, a population with mature hepatocyte-like features, and a hepatocyte-like population enriched in progenitor markers. This is compatible with the parenchymal cell heterogeneity described in patients with ALD due to the loss of mature hepatocyte identity or increased biological processes involved in biliary-to-hepatocyte transdifferentiation. Similarly, b-Org^HEP^ data show the presence of a biliary population expressing cholangiocyte markers but also pro-inflammatory features. This reactive cholangiocyte profile is typically found in ductular reaction cells in chronic liver disease when they proliferate and acquire reactive, pro-inflammatory and progenitor features[22,44,45]. In addition, we have used scRNA-seq to identify two clusters of cells with progenitor traits that differ in the level of expression of mature biliary/hepatocyte transcripts, thus indicating a continuum of states from progenitor to more mature cell populations. However, it is uncertain to what extent these progenitor cell populations are present in the human liver tissue. Interestingly, b-Org^HEP^ signatures are enriched in epithelial cell populations identified by scRNA-seq analysis of ALD liver tissue, thus suggesting that b-Orgs^HEP^ recapitulate liver disease epithelial heterogeneity. Altogether, we show that b-Orgs^HEP^ reproduce *in vitro* the epithelial compartment of the injured liver. However, b-Orgs still lack non-parenchymal cells. Therefore, efforts should be made to increase organoid complexity for improved disease modeling.

Here, we report proof of concept that b-Orgs^HEP^ retain ALD stage features and show the potential to identify molecular drivers of the disease. Notably, we found that b-Orgs^HEP^ from advanced stages are enriched in extracellular matrix remodeling, EMT and inflammation pathways, which are well-described processes involved in ALD progression. Moreover, we identified ELF3 as a potential driver of disease progression within the hepatocyte-like population of b- Orgs^HEP^. ELF3 has been implicated in the transcriptional reprogramming of hepatocytes in animal models of metabolic dysfunction-associated steatohepatitis (MASH), leading to loss of hepatocyte identity and the acquisition of biliary characteristics[42], as well as in the EMT process in the context of hepatocellular carcinoma[46]. Both hepatocyte dedifferentiation and EMT are hallmarks of advanced stages of ALD[41], but also of liver regeneration[47]. Indeed, Guo R et al. proved the association between ELF3 and inflammation-induced proliferation in cultured hepatocytes[48], suggesting a potential role in liver injury repair. Nonetheless, further investigation into the exact roles of ELF3 in chronic liver disease is required. These results support the relevance of using patient-derived liver samples as a source for organoid generation, which may preserve activated injury-related mechanisms alongside genetic and epigenetic modifications.

b-Orgs^HEP^ mimic some of the pathophysiological processes associated with ALD, including inflammation, ROS production and steatosis in response to ALD drivers. To reproduce ALD progression, we cultured b-Orgs^HEP^ with an AH-medium containing pro-inflammatory mediators and ethanol. Notably, the treatment with AH-medium induced morphological changes and decreased proliferation. According to the literature[49,50], our results may indicate that hepatocytes undergo metabolic adaptations leading to impaired energy flow that is translated into a dysfunctional, quiescent phenotype. Furthermore, b-Orgs^HEP^ responded to prednisolone treatment, the only approved drug therapy for AH, thus opening the door to drug testing applications. Altogether, we provide a patient-derived model for disease modeling and drug testing, thus overcoming current limitations in the development of new treatments for ALD, partially hampered by the lack of accurate animal models of advanced disease and relevant human-based pre- clinical models.

In this study we report on a methodology that enables the generation of patient-derived liver organoids recapitulating the heterogeneity of the hepatic epithelium. Moreover, we show the potential of b-Orgs to mimic liver epithelial plasticity and as a source for exploring molecular drivers of the disease. b-Orgs can be efficiently generated from different stages of chronic liver disease, providing a platform for patient-tailored disease modeling and drug testing.

## Abbreviations

ALD: alcohol-associated liver disease
ALT: alanine aminotransferase
AH: alcohol-associated hepatitis
AP: alkaline phosphatase
AST: aspartate aminotransferase
BEC: biliary
b-Orgs: biopsy-derived organoids
Chol-Orgs: cholangiocyte organoids
CLF: cholyl-L-lysyl-fluorescein
DEGs: differentially expressed genes
EMT: epithelial-to-mesenchymal transition
EtOH: ethanol
GSEA: gene set enrichment analysis
HB: hepatobiliary
HEP: hepatocyte
IL1b: interleukin-1 beta
INR: international normalized ratio
LDH: lactate dehydrogenase
LiGEP: liver-specific gene expression panel
LPS: lipopolysaccharide
NES: normalized enrichment score
OPN: osteopontin
PAMPs: pathogen-associated molecular patterns
ROS: reactive oxygen species
scRNA-seq: single-cell RNA sequencing
TNFa: tumor necrosis factor alpha
UMAP: uniform manifold approximation and projection
WGCNA: weighted correlation network analysis.

## Acknowledgments

This work has been developed at the Centre Esther Koplowitz (CEK). The authors want to thank Pepa Ros for her excellent laboratory management support. We thank Xavier Thillen for his experimental contribution. We are indebted to the Biobank Unit and the cytometry and cell sorting facility of the Institut d’Investigacions Biomèdiques August Pi i Sunyer (IDIBAPS) for their technical help. We thank the TEM-SEM Electron Microscopy Unit from the Scientific and Technological Centers (CCiTUB), of the University of Barcelona, and staff for their support and advice on TEM technique.

## Author contributions

S.A. participated in the design of the study, performed most of the experiments and drafted the manuscript; L.Z. and R.F-L. contributed to organoid experiments and critically reviewed the manuscript. R.A. M-G., B.A- B., P.C-V., L.S-V. and Z.X. assisted with cell culture and critically reviewed the manuscript. JJ.L. and L.Z. performed bulk RNA sequencing analysis. R.A. M-G., G.S. and AR participated in organoid scRNA-seq analysis and reviewed the manuscript. E.P. and J.G-G. recruited patients for the study and contributed to the critical review of the manuscript. A.B-R. and M.P. coordinated biopsy collection. PR-B contributed to organoid histological characterization. S.A. generated scRNA-seq data from ALD biopsies. M.C., I.O., S.A., A.M., E.M. and R.B. interpreted data and contributed to the critical review of the manuscript. E.P. recruited patients for the study and critically reviewed the manuscript. P.S-B. conceived and designed the study, critically reviewed the manuscript and supervised and coordinated the study.

