## Supplemental Data 1 for "Patient-derived liver biopsy organoids enable precision alcohol-associated liver disease modeling"

**The PDF file includes:**

Materials and Methods

Figs and legends. S1 to S6

Table S1

Legend for Table S2

Legends for movies

References

**Other Supplementary Material for this manuscript includes the following:**

Table S2

Movies S1 to S2

### SUPPLEMENTARY MATHERIALS

#### Biopsy culture and organoid generation and expansion

Fresh patient-derived needle-biopsies were obtained at early (n= 28) and advanced (n=34) stages of ALD. Liver biopsy fragments around 2-3mm were cut using a scalpel and plated in 50uL-drops of Matrigel (Corning) in 24 well-plates. After Matrigel was solidified, hepatocyte (HEP) or biliary (BEC) mediums were added.

HEP-medium is prepared as described by Hu H et al.[1] and consists of Advanced DMEM/F12 (Thermo Fisher Scientific, 12634-010) supplemented with 1% HEPES (Gibco, 15630-056), 1% GlutaMax (Gibco, 35050-038), 1% Pen-Strep (Biowest, L0022), 1x N2 supplement (Gibco, 11520536), 1x B27 supplement (without vitamin A) (Gibco, 12587010), 500ng/mL recombinant human (rh) RSPO1 (Stemcell Technologies, 78213), 50ng/mL rhHGF (GenScript, Z03229), 100ng/ml rhFGF10 (GenScript, Z03314), 50ng/mL rhEGF (GenScript, Z00333), 100ng/mL rhFGF7 (GenScript, Z03047), 20ng/mL rhTGFa (GenScript, Z03332), 1.25mM N-acetylcysteine (Sigma, A9165), 10nM human [Leu15]-gastrin I (Sigma, G9145), 10mM Nicotinamide (Sigma, N0636), 3µM CHIR99021 (Tocris, 1386) and 2µM A8301 (Tocris, 1421).

BEC-medium is prepared as described by Broutier L et al.[2] with minor modifications and consists of Advanced DMEM/F12 (Thermo Fisher Scientific, 12634-010) supplemented with 1% HEPES (Gibco, 15630-056), 1% GlutaMax (Gibco, 35050-038), 1% Pen-Strep (Biowest, L0022), 1x N2 supplement (Gibco, 11520536), 1x B27 supplement (without vitamin A) (Gibco, 12587010), 500ng/mL recombinant human (rh) RSPO1 (Stemcell Technologies, 78213), 25ng/mL rhHGF (GenScript, Z03229), 100ng/ml rhFGF10 (GenScript, Z03314), 50ng/mL rhEGF (GenScript, Z00333), 125ng/mL rhNoggin (Preprotech, 120-10C), 1.25mM N-acetylcysteine (Sigma, A9165), 10nM human [Leu15]-gastrin I (Sigma, G9145), 10mM Nicotinamide (Sigma, N0636) and 5µM A8301 (Tocris, 1421), 0.5 µM CHIR99021 (Tocris, 1386) and 10µM Forskolin (Tocris, 2264).

Medium was changed every 2 days and 10µM Rho Inhibitor γ-27632 (Axon Medchem, 1683) was added to the medium during the first week of fine-needle biopsies culture. After 12-18 days, b-Orgs were mechanically detached from the biopsy and transferred to a 15mL-tube to be passaged to 3-4 wells following Broutier et al. protocol[2] with minor modifications. Grown b-Orgs can be subsequently passaged 1:2 – 1:3 until passage 8 every 7-15 days or cryopreserved in Cell Recovery Freezing Medium (Gibco).

After separation of budding organoids from biopsies, biopsies were frozen in Cell Recovery Freezing Medium (Gibco) or transferred to a new 50uL-Matrigel drop to provide a subsequent generation of b-Orgs as previously described.

Standard organoid lines from liver resections were established and expanded with minor modification following Broutier et al. protocol[2] and using BEC-medium.

Organoids morphology and growth were tracked by bright field imaging using Olympus IX51 inverted microscope. Time lapse showing organoid growth from biopsies was performed by taking bright field images using tile scan option every 12 hours for 3 days. Leica AF6000 microscope was used for mosaic images.

All organoid lines used in this study has been confirmed to be negative for mycoplasma by PCR.

### Organoid staining

#### *Organoid immunofluorescence*

Organoid immunofluorescence was performed using whole organoids or with organoids paraffin sections. Organoids were stained for EPCAM (1:100, Dako, M0804), ALB (1:200, Sigma, A7544), HNF4A (1:200, abcam, ab92378), TROP2 (1:50, R&D, AF1122), MRP2 (1:100, abcam, ab3373) and CDH1 (1:100, Cell Signaling, 3195), SOX9 (1:500, Millipore, ab5535), KRT7 (1:50, Dako, M701801), GS (1:100, Abcam, ab73593), NCAM (1:50, Sigma, C9672), Osteopontin (1:100, Abcam, ab8448), Vimentin (1:100, Leica, NCL-VM-V9), and PDGFRb (1:100, abcam, ab32570).

For paraffined organoids stainings, organoids were first fixed with 4% PFA within the Matrigel drop. Then, organoids were washed and the whole drop containing the organoids was included in HistoGel (FisherScientific, 12006679) following the manufacturers' protocol and dehydrated before being included in paraffin. Organoid paraffin sections (5um) were deparaffinized and incubated in Target Retrieval Solution (Citrate Ph6, DAKO, Glostrup, Denmark), heated in a pressure cooker for 20 minutes, or rehydrated and antigen retrieved with EnVision Flex Target Retrieval Solution Low or High PH in a DAKO PT Link before adding primary antibodies.

For whole organoid staining, organoids were removed from Matrigel by incubating on ice for 1-2 hours with mild shaking using Cell Recovery Solution (Corning, 354253). After washing, organoids were pelleted at 75g for 2 minutes, fixed with 4% PFA and transferred onto a slide using Thermo Scientific™ Cytospin™ 4 Cytocentrifuge. PFA was quenched by 20mM glycine rinse and organoids were washed and permeabilized for 30 minutes at room temperature with 0.2% Triton X-100 in PBS. Next, organoids were washed with washing solution (0.2% Triton X-100, 0.02% Tween 20 in PBS) and blocked with 1% BSA in washing solution for 45 minutes at room temperature before labelling with primary antibodies.

After performing antigen retrieval on sections and blocking whole organoids, both sections and whole organoids were incubated overnight at 4ºC with primary antibodies. After washing in PBS, sections and whole organoids were incubated with secondary antibodies donkey anti-rabbit Cy3 (1:400, Jackson ImmunoResearch, 111-165-003), donkey anti-goat Cy3 (1:200, Jackson ImmunoResearch, 705-165-147), donkey anti-rabbit Alexa Fluor 488 (1:200, Jackson ImmunoResearch, 711-545-152), and goat anti-mouse Alexa Fluor 488 (1:400, Invitrogen, A11001) for 30 minutes at room temperature and eventually mounted with Mounting Medium for Fluorescence with 4',6-diamidino-2-phenylindole (DAPI) (Vector Laboratories, Burlingame, CA) for nuclear staining.

*Glycogen and H&E stainings*

Organoids paraffin sections (5um) were used for standard H&E and PAS staining protocols.

For H&E and PAS stainings, sections were dewaxed with xylol, hydrated through serial ethanol dilutions and finally rinsed in water. In the case of H&E, tissue sections were stained with hematoxylin (Sigma-Aldrich) for 10 minutes and eosin for additional 5 minutes. For PAS staining, sections were incubated with periodic acid solution for 5 minutes, Schiff's reagent for 15 minutes and nuclei were counterstained with hematoxylin. In both cases, residual dyes were washed, and sections were dehydrated and mounted in DPX medium (Merck).

**RNA Isolation and mRNA expression analysis**

Total RNA was extracted from total liver tissue and from cultured organoids using the commercial RNeasy Micro and Mini Kits (Qiagen) following manufacturer's instructions. Total RNA extracted was assessed by Nanodrop (ThermoScientific). Extracted RNA was reverse-transcribed with the High-Capacity cDNA Reverse Transcription kit (Applied Biosystems; Carlsbad, CA). Gene expression quantification was accomplished using Taqman gene expression assay probe and primers, and Platinum SYBR Green qPCR SuperMix (Thermofisher, Ref. 11733046) and PrimeTime® qPCR Primers (IDT). Quantitative real-time PCR (qPCR) was performed with an ABI 7900 HT cycler (Life Technologies) and QuantStudio 7 Pro (Applied Biosystems). Expression was normalized to the expression of the *18S* or *GAPDH* housekeeping genes. Expression values were calculated based on the ∆∆Ct method. The results were expressed as 2^-∆∆Ct.

**Bulk RNA-Sequencing**

*Library preparation and RNA sequencing*

The quantity and quality of the RNAs were evaluated using Qubit RNA HS Assay Kit (Thermo Fischer Scientific, Cat#Q32855) and Agilent RNA 6000 Nano Chips (Agilent Technologies, Cat #5067-1511), respectively. Sequencing libraries were prepared using NuGEN Universal Plus mRNA-Seq kit (PART NO. 0508, 9133, 9134) and following the user guides (M01442v2 and M01485v8). Then mRNA was purified, fragmented and transcribed to cDNA. Afterwards, end-repair and adaptor ligation were carried out. Finally, after strand selection, enrichment of libraries was achieved by PCR. Before and after purification, libraries were visualized on an Agilent 2100 Bioanalyzer using Agilent High Sensitivity DNA kit (Agilent Technologies, Cat. #5067-4626) and quantified using Qubit dsDNA HS DNA Kit (Thermo Fisher Scientific, Cat. #Q32854). Sequencing using 101-bp paired-end reads was performed using NovaSeq6000 Illumina platform. The RNA sequencing experiments were performed at the Genomics Unit at CICbioGUNE.

*RNA sequencing analysis*

Raw reads were analyzed for data quality and filtered using skewer*[3]* for removing the low-quality reads and trimming the Illumina adapter. STAR program*[4]* against Homo Sapiens (GRCh38) was used for mapping the reads followed by the quantification of genes with the RSEM program*[5]* using GENCODE v26 reference annotation. We used TMM method and limma-voom transformation to normalize the non-biological variability. Differential expression between different groups was assessed using moderated t-statistics*[6]* after selecting no autosomal protein-coding genes with at least a value of 10 reads in a sample.

From normalised expression matrix we computed corresponding Z-scores. Principal component plots were performed using R statistical software.

Liver tissue similarity (LiGEP algorithm) was calculated using organoids gene expression in FPKMs as an input data set for the interface named Web-based Similarity Analytics System (W-SAS)*[7,8]*. Heatmaps were performed using z-score values and the package pheatmap v1.0.12 in R v4.3.0. Volcano plot was created using SR plot software.

GSEA software v4.2.2 (Broad Institute) was used in pre-ranked list mode to assess the following gene enrichment analysis. Cell type signatures were obtained from scRNA-seq data sets available in the literature (biliary*[9–12]*, adult/fetal hepatocyte*[9,11–13]*, mesenchymal*[9]* ) populations and used to explore enrichment when comparing a) standard organoids with b-orgs or b) b-Orgs$^{HEP}$ with b-Orgs$^{BEC}$. Genes from organoids comparisons were pre-ranked by t-statistics parameter. For studying b-Org$^{HEP}$ resemblance to their parental tissue, top 500 DEGs for early and advanced biopsies were considered from their direct comparison (filtered as FC > 2 or <-2 and adjusted p-value < 0.05 and rasnked by FC). Both signatures were used for enrichment analysis when comparing b-Orgs$^{HEP}$ from early stages versus advanced stages. Genes from organoids comparison were pre-ranked by t-statistics parameter. Core enrichment genes for both disease stages were used for further enrichment analysis using EnrichR software for biological processes (Reactome, Kegg and Gene Ontology databases were considered) and Chea3 for upstream regulator activity prediction. To study potential changes in organoid cell populations when stimulated with AH-medium, DEGs from each cluster resulting from scRNA-seq analysis were used as an input gene list for GSEA enrichment. In the case that more than 500 DEGs characterised a cluster, top 500 genes ordered by FC were considered. As previously mentioned, comparison of vehicle and AH-medium stimulated organoids were ranked by t-statistic.

To carry out WGCNA*[14]* we choose a group of 174 genes that exhibits difference between any of 2 sets of groups (p-value < 0.05 and FC > 2). The PickSoftThreshold function was used to select the soft threshold power used to construct a network based on the criterion of approximate scale-free topology.

We choose the power 13.5, which is the lowest power for which the scale-free topology fit index curve flattens out upon reaching RsquaredCut a value of 0.8. The adjacency matrix was calculated and was converted into a topological overlap matrix (TOM), and the degree of dissimilarity between genes was calculated and a full dendogram were built. Gene modules were generated using the cutreeDynamic function setting the minimum number of genes as 20 and a higher sensitivity to cluster splitting controlled by the deepSplit parameter (deepSlipt=1). Finally, modules with a dissimilarity coefficient less than 0.2 were merged. We computed the eigengene for each module, defined as the first principal component of the module representing the overall expression level of the module. Next, to reduce multi-dimensionality, eigengenes for each module were used to relate the modules to different set of groups (b-Org generation comparison and b-Org passages comparison) phenotypes by means of wilcoxon rank sum test computation measures. Finally, we apply enrichment analysis using for each of the 3 modules obtained using Gene Ontology using clusterProfiler package*[15]*.

**Single cell RNA sequencing (scRNA-seq)**

*Sample preparation, library construction and sequencing*

b-Orgs$^{HEP}$ (n = 4; n=1 early stage, n= 3 advanced stage) were dissociated into single cell by incubating for 30 minutes in a water bath with gently shaking at 37ºC with TrypLE Express (Gibco, 12604021) supplemented with 2mM EDTA (Life Technologies, 15575-020). Next, mild pipetting was performed to ensure organoids disaggregation. Cell suspension was filtered using a 40-um nylon mesh, pelleted, resuspended in 0.04% BSA in DPBS-/- and transported on ice until further processing.

Liver biopsies from patients with ALD (n=2) were processed and dissociated into single cell suspensions following the protocol reported by MacParland SA et al.*[12]*.

*scRNA-seq processing and analysis*

Single cell suspensions, with a concentration of 700-1200 cells/μl, were loaded into the Chromium Next GEM Chip G wells, to have a final recovery of approximately 5.000 cells per suspension. The Chromium Controller performed single cell partitioning and barcoding. Cells were partitioned into nanoliter-scale Gel Beads-in-emulsion (GEMs), where all generated cDNA share a common 10x Barcode. A pool of ~3,500,000 10x Barcodes were then sampled separately to index each cell's transcriptome. Dual Indexed libraries generated and sequenced from the cDNA and 10x Barcodes were used to associate individual reads back to the individual partitions. Single Cell 3′ v2 libraries were generated following the Single Cell 3′ v2 Reagent Kits User Guide (Document CG00052, 10X genomics). The obtained libraries were sequenced on the Illumina NovaSeq 6000 platform at 40,000 reads/cell with PE-150 strategy.

We performed sequence alignment of our sample reads, sourced from FASTQ files, against the GRCh38 reference genome and conducted pre-processing using the Cell Ranger count function provided by 10X Genomics (v7.1.0) streamlining the data for subsequent downstream analyses.

A filtered gene expression matrix containing the number of UMI for every cell and gene was created for each sample. A standard pre-processing workflow for scRNA-seq data was performed using R v4.3.0 and the package Seurat v4.3.0 (Butler et al., 2018; Stuart et al., 2019).

Organoids raw data was merged in a unique seurat object, resulting in 36,601 genes and 21,999 cells. Cells presenting less than 1,000 unique genes or/and more than 15% of mitochondrial genes were discarded. After quality control, the data input contained 36,601 genes and 14,183 cells and was normalized using the Seurat function NormalizeData with the global-scaling normalization method "LogNormalize". Genes with high cell-to-cell variation were identified by FindVariableFeatures function. A total of 2,000 genes were returned as top variable features and used to generate the PCA dimensionality reduction after scaling data with the ScaleData function. To correct batch effects, Harmony v.0.1.1 was used (ref) together with the FindNeighbors function of Seurat, considering "harmony" reduction and 20 dimensions. Finally, Louvain algorithm implemented in the FindClusters function was applied to cluster the harmonized data. A clustering resolution of 0.4 of the batch-corrected data set was used in all the following analysis. However, after annotation, cluster 0 and 10 were integrated as the "Progenitor" cluster due to similarity between both populations resulting in a total number of 11 clusters. Clusters were visualized using Uniform Manifold Approximation and Projection for Dimension Reduction (UMAP). Differentially expressed genes (DEGs) were calculated for each cluster considering a) genes that showed at least 0.25-fold change (log-scale) on average between the two populations and b) genes detected in a minimum percentage of 25% of the cells in either of the two groups of cells. Gene expression levels and density were visualized using DotPlot function in Seurat and UMAP reduction in Nebulosa v.1.10.0, respectively. Cell cycle scores were calculated using CellCycleScoring function in Seurat. Pathway functional enrichment was performed using DEGs with an adjusted p-value < 0.05 for each cluster as an input data set for EnrichR*[16]*. Cluster annotations were performed taking into consideration canonical markers of epithelial (biliary, hepatocyte, progenitor)*[9,11,12,17]* and mesenchymal cell populations*[9]* and enriched biological processes associated to each cluster. Gene regulatory network activity was inferred following SimiC pipeline as described previously*[18]*.

ALD liver tissue raw data was merged in a unique Seurat object. Cells presenting less than 300 features or more than 3,000 features were discarded. Additionally, only cells with less than 50% of mitochondrial genes were analysed. After quality control, 6,659 cells and 33,538 genes were analysed following the same pipeline as previously described for organoids data. Again, batch effects were corrected using Harmony v.0.1.1 (ref) together with the FindNeighbors function of Seurat, considering “harmony” reduction and 30 dimensions. A clustering resolution of 0.5 determined by Louvain algorithm and DEGs calculated for each cluster as previously stated for organoids were used to annotate cell populations based on the expression of canonical markers*[19]*. From the resulting 20 clusters, 3 of them were annotated as epithelial cells and subseted into a new Seurat object containing 753 cells. Data from the epithelial compartment was then normalized, cell-to-cell variation were identified, PCA dimensionality reduction was calculated and data was scaled following the same pipeline as before. FindNeighbors function of Seurat considering PCA reduction and 20 dimensions was run and Louvain algorithm in FindClusters function was used to subcluster the epithelial fraction. Cluster were visualized uding UMAP and DEGs from a resolution of 0.5 were calculated with the FindAllMarkers function as stated for organoids and used to label cell populations. Finally, signatures from each non-cycling epithelial organoid cluster consisting in the 25 genes with higher FC were used to calculate module scores in each of the epithelial liver tissue clusters were using AddModuleScore from Seurat and visualized by DotPlot function.

**Correlation analysis**

To study ELF3 clinical relevance, we used the data set published in 42 containing total liver tissue transcriptomic data and clinical information from an ALD cohort*[21]*. Correlations between variables were evaluated using Spearman's rho or Pearson's r, when appropriate.

**Transmission electron microscopy (TEM)**

Organoids were harvested from Matrigel as stated in the Organoids staining section. Once in suspension, organoids were fixed with 2.5 % glutaraldehyde and 2% paraformaldehyde in 0.1 M phosphate buffer. Samples were post fixated with osmium tetroxide and dehydrated with acetone, embedded in Spurr resin and sectioned using Leica ultramicrotome UC7 (Leica Microsystems). Ultrathin sections (50–70 nm) were stained with 2 % uranyl acetate for 10 min, a lead-staining solution for 5 min and then analysed with a transmission electron microscope, JEOL JEM-1010 fitted with a Gatan Orius SC1000 (model 832) digital camera at the TEM-SEM Electron Microscopy Unit, Scientific and Technological Centers of the University of Barcelona (CCiTUB).

**Albumin quantification**

Organoids medium was collected at 72 hours after the medium change when 80-95% confluence was reached. Albumin levels were analysed using the Human Albumin ELISA Kit (Bethyl Laboratories; E80-129) and normalized by cell number at the time of medium collection.

**Cholyl-Lys-Fluorescein (CLF) incorporation assay**

Organoids were removed from Matrigel by incubating on ice for 2 hours with mild shaking using Cell Recovery Solution (Corning, 354253). Next, organoids were washed, resuspended in HEP-medium and incubated in suspension for 1 hour at 37ºC in 5% $CO_2$ with 5uM CLF (Apollo Scientific, #54-BICR355). Following, organoids were washed three times with DPBS-/- and nuclei were counterstained with Hoechst 33342 (Invitrogen, H3570). Finally, CLF incorporation was visualized using Olympus IX51 microscope.

**Disease progression modelling and prednisolone treatment**

To model disease progression, b-Orgs$^{HEP}$ were treated in duplicate for 5 days with AH-medium or ethanol (EtOH). AH-medium consisted of HEP-medium with 100ng/mL LPS (Sigma, L2637), 20ng/mL rhTNFa (Genscript, Z01001), 10ng/mL rhIL1b (Preprotech, 200-01B) and 100mM EtOH. Control group was just cultured with HEP-medium. Control medium and whole AH-medium were changed every 2 days and ethanol was daily refreshed. To limit EtOH evaporation, parafilm was used to seal the plate. To test b-Orgs$^{HEP}$ response to corticoids, prednisolone (Sigma, P6004) was daily added to the AH-medium or control HEP-medium at 10, 20 and 50uM during the last three days of treatment. After the 5-day treatment, b-Orgs$^{HEP}$ were taken in RLT for RNA extraction or processed for further analysis as described as follows.

*Proliferation analysis*

To assess organoid proliferation after treatment with AH-medium, b-Orgs$^{HEP}$ were harvested from Matrigel, fixed, and stained for KI67 (1:50, abcam, ab16667) and EPCAM (1:100, Dako, M0804) as explained in the Organoids staining section.

Secondary antibodies donkey anti-rabbit Cy3 (1:400, Jackson ImmunoResearch, 111-165-003) and goat anti-mouse Alexa Fluor 488 (1:400, Invitrogen, A11001) were used for immunofluorescence labelling. Images for quantification were acquired using Leica DM2500 confocal microscope. Percentage of KI67+ EpCAM+ nuclei versus total nuclei was quantified in 8 images per condition by Fiji Software.

Alternatively, ATP levels were measured after AH-medium and prednisolone treatment in white 96-well plates (Greiner, 655088) following CellTiter-Glo® 3D Cell Viability Assay (Promega) manufacturer's protocol. Luminescence was measured with Tecan Infinite MNano+ 200 Pro.

*Lipid accumulation staining*

b-Orgs$^{HEP}$ were plated in 48-well plates (Costar, 3548) and treated in duplicates. After treatment, organoids were harvested from Matrigel with Cell Recovery Solution (Corning, 354253) as explained with further details in the Organoids staining section. After fixation and transference onto a slide using Thermo Scientific™ Cytospin™ 4 Cytocentrifuge, b-Orgs$^{HEP}$ were stained for Oil-red-O (Sigma, 01391). Briefly, organoids were washed with distilled water and incubated with 60% isopropanol for 5 minutes. Next, b-Orgs$^{HEP}$ were incubated with 60% Oil-red-O reagent diluted in distilled water for 40 minutes at room temperature. Finally, nuclei were counterstained with haematoxylin after washing and mounted in Aquatex (Merck, 108562). Images for quantification were acquired using Olympus IX51 microscope. Oil-red-O staining area was quantified and normalized by total organoid area using Fiji software.

*Cell death and apoptosis assessment*

After treating b-Orgs$^{HEP}$ with AH-medium in 96-well plates (Costar, Corning), b-Orgs$^{HEP}$ were first dissociated into single cell by incubating for 30 minutes in a water bath at 37ºC with TrypLE Express (Gibco, 12604021) supplemented with 2mM EDTA (Life Technologies, 15575-020). First, cells were stained with anti-EpCAM-eFluor660 (1:100, eBioscience, 50-9326-42). Then, FITC Annexin V (Biolegend, 640922) was used to assess cell death and apoptosis following the manufacturing protocol with minor modification. Briefly, cells were washed with Annexin V Binding Buffer and incubated for 15 minutes at room temperature with 5uL of FITC Annexin V. Then, the excess of Annexin V probe was washed with Annexin V Binding Buffer and finally, cells were resuspended in Annexin V Binding Buffer and 7AAD was added before analysing with BD FACS Canto III and further analysed by FlowJo software.

*ROS index determination*

b-Orgs$^{HEP}$ plated and treated in 96-well plates (Costar, Corning) were used for ROS measurement. CellROX Deep Red reagent (Invitrogen, C10422) was added at a concentration of 5uM and incubated at 37ºC for 30 minutes. Then, b-Orgs$^{HEP}$ were dissociated into single cell as explained in the previous section (Cell death and apoptosis assessment) and viable cells were further analysed for ROS detection using BD FACS Canto III and further analysed by FlowJo software. ROS index was calculated as the percentage of ROS+ alive cells multiplied by ROS median intensity of alive cells.

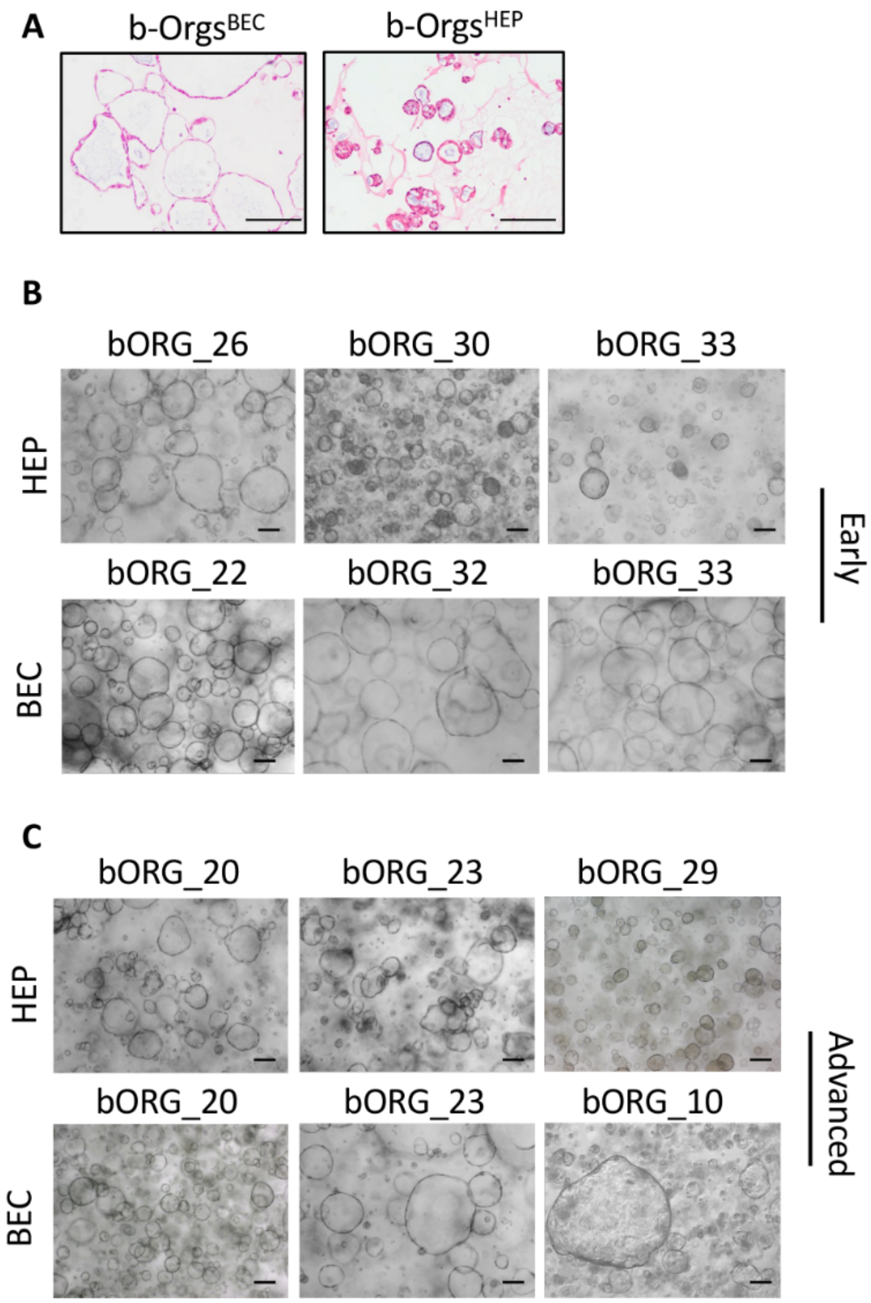


**Supplementary Figure 1. Derivation of liver organoids from needle-biopsies at different stages of ALD. Related to Figure 1.**

**(A)** Representative H&E staining images of b-Orgs cultured with BEC and HEP culture conditions. Scale bar = 50 μm. **(B)** Representative bright field images of b-Orgs from early stages of ALD using both HEP and BEC culture mediums. Scale bar = 200 μm. **(C)** Representative bright field images of b-Orgs from advanced stages of ALD using both HEP and BEC culture mediums. Scale bar = 200 μm. ALD, alcohol-associated liver disease; b-Orgs, biopsy-derived organoids.

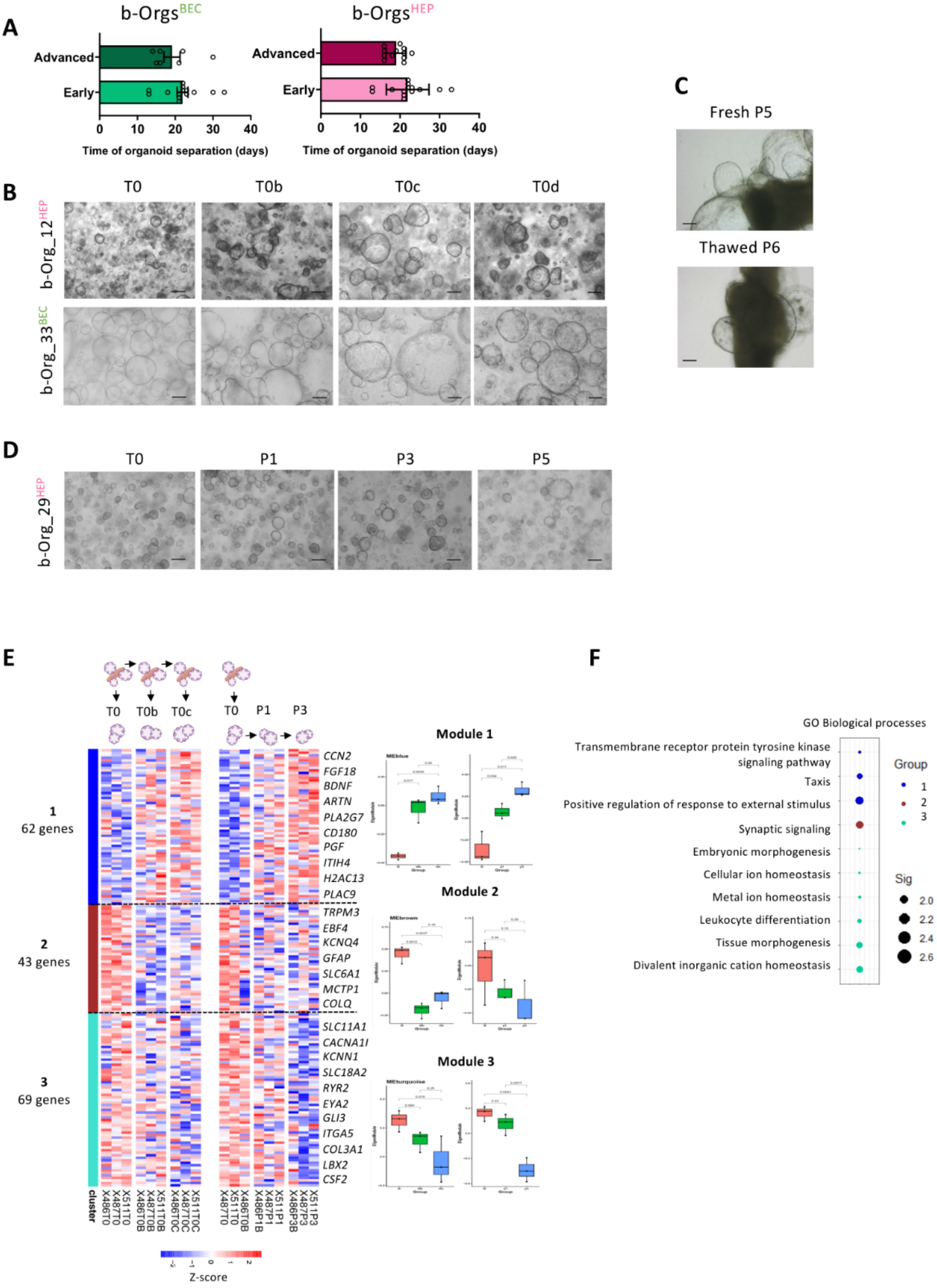

**Supplementary Figure 2. Biopsies can be maintained long-term in culture while generating organoids. Related to Figure 2.**

**(A)** Graphs showing the number of days needed to separate organoids from the biopsy, starting to count from the day that the biopsy was embedded in Matrigel. Data is presented as mean ± SEM. No significant differences were found as assessed by unpaired t test. **(B)** Representative bright field images of successive b-Org generations using both BEC and HEP culture conditions. Scale bar = 200 µm. **(C)** Representative bright field image of fresh and thawed biopsy with budding b-Orgs$^{BEC}$. Scale bar = 200 µm. **(D)** Representative bright field image of the first (T0) generation of b-Orgs$^{HEP}$ being further expanded up to passage (P) 5. Scale bar = 200 µm. **(E)** Heatmap and boxplots depicting gene modules obtained by WGCNA analysis that significantly change across b-Org generations and passages (P) (n=3 per group). **(F)** Significantly enriched GO biological processes for each of the three WGCNA gene modules. Dot plot size represents -log10(p.adjusted) values and the colour refers to the gene module. ALD, alcohol-associated liver disease; b-Orgs, biopsy-derived organoids.

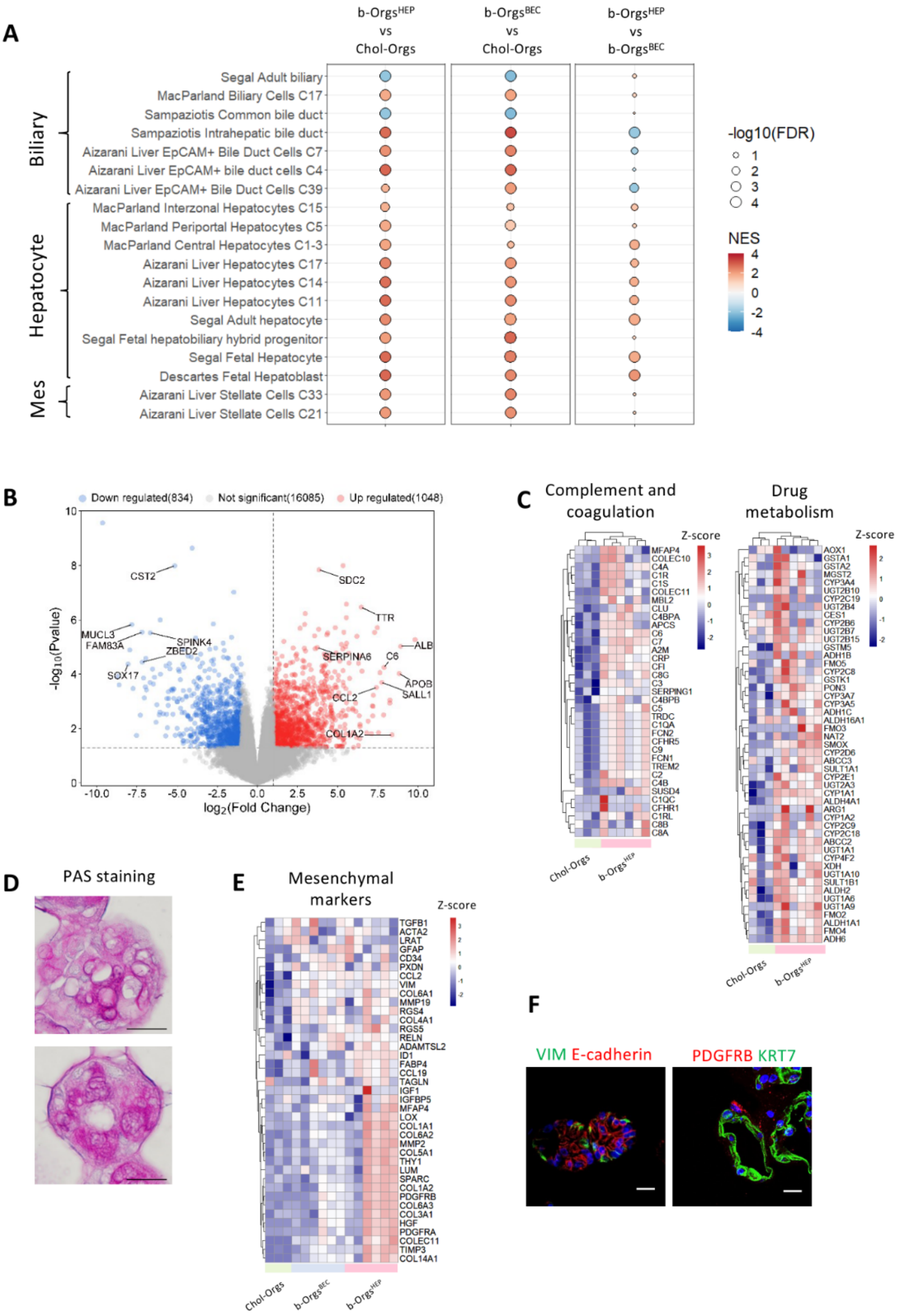
A
b-Orgs^HEP vs Chol-Orgs
b-Orgs^BEC vs Chol-Orgs
b-Orgs^HEP vs b-Orgs^BEC
Biliary
Hepatocyte
Mes
Segal Adult biliary
MacParland Biliary Cells C17
Sampaziotis Common bile duct
Sampaziotis Intrahepatic bile duct
Aizarani Liver EpCAM+ Bile Duct Cells C7
Aizarani Liver EpCAM+ bile duct cells C4
Aizarani Liver EpCAM+ Bile Duct Cells C39
MacParland Interzonal Hepatocytes C15
MacParland Periportal Hepatocytes C5
MacParland Central Hepatocytes C1-3
Aizarani Liver Hepatocytes C17
Aizarani Liver Hepatocytes C14
Aizarani Liver Hepatocytes C11
Segal Adult hepatocyte
Segal Fetal hepatobiliary hybrid progenitor
Segal Fetal Hepatocyte
Descartes Fetal Hepatoblast
Aizarani Liver Stellate Cells C33
Aizarani Liver Stellate Cells C21
-log10(FDR)
1
2
3
4
NES
4
2
0
-2
-4
B
Down regulated(834)
Not significant(16085)
Up regulated(1048)
-log10(Pvalue)
log2(Fold Change)
CST2
MUCL3
FAM83A
SPINK4
ZBED2
SOX17
SDC2
TTR
ALB
SERPINA6
C6
APOB
SALL1
CCL2
COL1A2
C
Complement and coagulation
Drug metabolism
Z-score
Chol-Orgs
b-Orgs^HEP
D
PAS staining
E
Mesenchymal markers
Z-score
TGFB1
ACTA2
LRAT
GFAP
CD34
PXDN
CCL2
VIM
COL6A1
MMP19
RGS4
COL4A1
RGS5
RELN
ADAMTSL2
ID1
FABP4
CCL19
TAGLN
IGF1
IGFBP5
MFAP4
LOX
COL1A1
COL6A2
MMP2
COL5A1
THY1
LUM
SPARC
COL1A2
PDGFRB
COL6A3
COL3A1
HGF
PDGFRA
COLEC11
TIMP3
COL14A1
Chol-Orgs
b-Orgs^BEC
b-Orgs^HEP
F
VIM E-cadherin
PDGFRB KRT7

**Supplementary Figure 3. b-Orgs present enriched liver-like and hepatobiliary features. Related to Figure 3.**

**(A)** Dot plot showing enrichment of liver cell type signatures in b-Orgs (n=6, BEC; n=6 HEP) when compared to Chol-Orgs (n=3) (two first columns) and in b-Orgs$^{HEP}$ when compared to b-Orgs$^{BEC}$ (third column). NES is shown as a scale colour and dot size represents significance. **(B)** Volcano plot comparing b-Orgs$^{HEP}$ (n=6) to Chol-Orgs (n=3). Genes coloured in red are significantly increased in the b-Org$^{HEP}$ group and genes coloured blue are down in the b-Org$^{HEP}$ group. **(C)** Heatmaps displaying scaled expression levels of genes involved in hepatocyte-related processes (Drug metabolism and coagulation/complement) in Chol-Orgs and b-Orgs$^{HEP}$. **(D)** Representative image of PAS staining in b-Orgs$^{HEP}$. Scale bar = 100 µm. **(E)** Heatmaps depicting scaled expression levels of mesenchymal genes in Chol-Orgs and b-Orgs. **(F)** Representative staining of mesenchymal (VIM and PDGFRB) and epithelial (E-cadherin, KRT7) markers in b-Orgs$^{HEP}$. Scale bar = 20 µm. b-Orgs, biopsy-derived organoids; Chol-Orgs, cholangiocyte organoids; GSEA, gene set enrichment analysis; NES, normalized enrichment score; PAS, periodic acid-Schiff; PCA, principal component analysis.

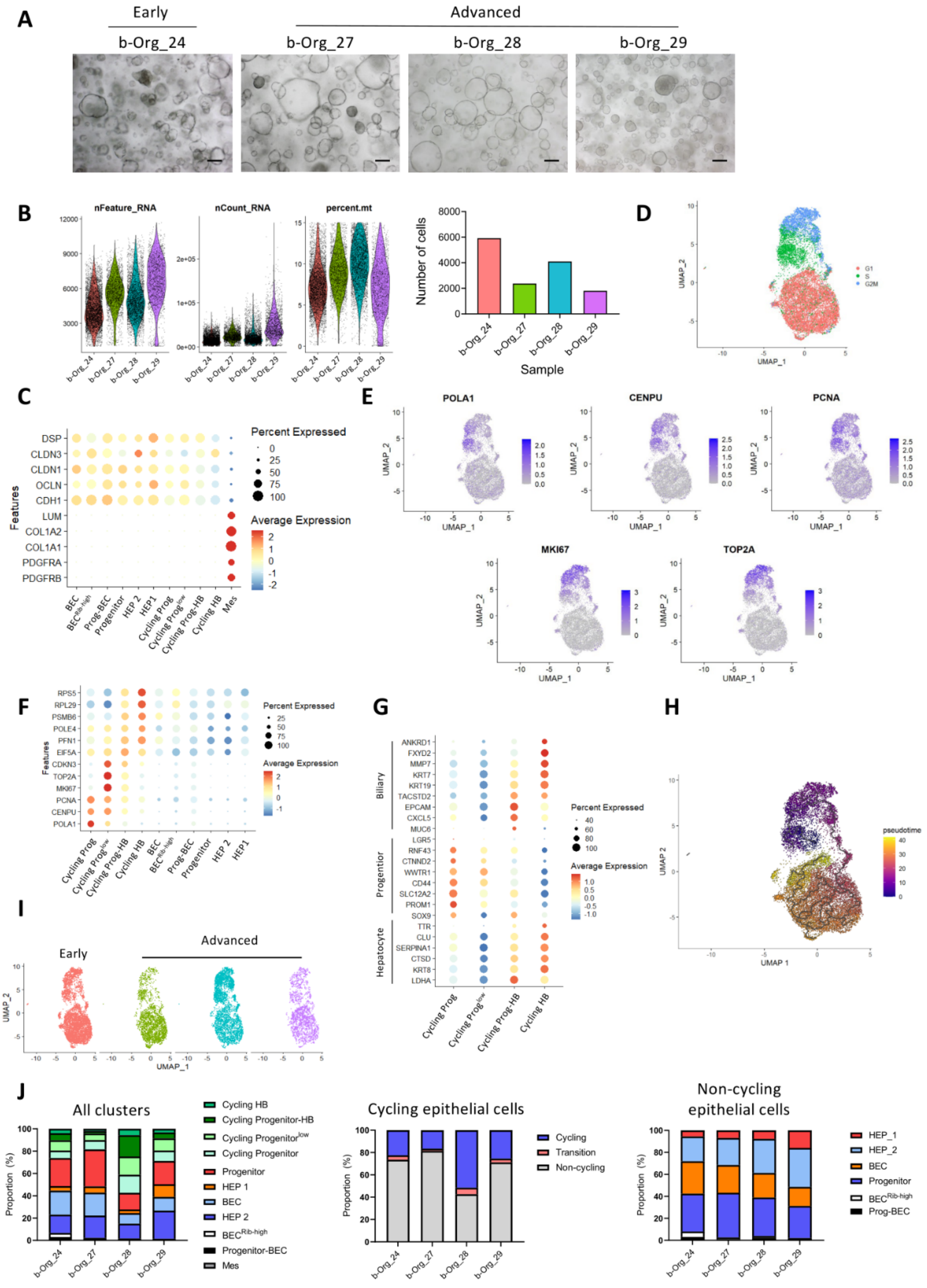


**Supplementary Figure 4. b-Orgs$^{HEP}$ recapitulate the liver epithelial heterogeneity. Related to Figure 4.**

**(A)** Representative bright field images of b-Orgs$^{HEP}$ used for scRNA-seq analysis (n=4). Scale bar = 200 µm. **(B)** Violin plots showing quality control filtering for each sample (left) and graph showing the resulting number of cells post-filtering that were used for downstream analysis (right). **(C)** Scaled average expression (dot colour) of epithelial and mesenchymal genes in each cluster. Dot size displays the percentage of cells within the cluster expressing the gene. **(D)** UMAP displaying proliferative cells (S and G2M groups) and non-cycling cells (G1) as assessed by CellCycleScoring function in Seurat. **(E)** UMAPs showing expression levels of cell cycle markers. **(F)** Scaled average expression (dot colour) of cell cycle-related genes in each of the epithelial populations. Dot size displays the percentage of cells within the cluster expressing the gene. **(G)** Scaled average expression (dot colour) of biliary, hepatocyte and progenitor markers in cycling clusters. Dot size displays the percentage of cells within the cluster expressing the gene. **(H)** UMAP showing pseudotime trajectories. **(I)** UMAPs of each of the analysed samples after harmonization. **(J)** Graphs showing for each sample the proportion in percentage (%): of every cluster, of cells in each cycling stage, and of each epithelial cell population. b-orgs, biopsy-derived organoids. b-Orgs, biopsy-derived organoids; UMAP, Uniform Manifold Approximation and Projection; scRNA-seq, single-cell RNA-sequencing.

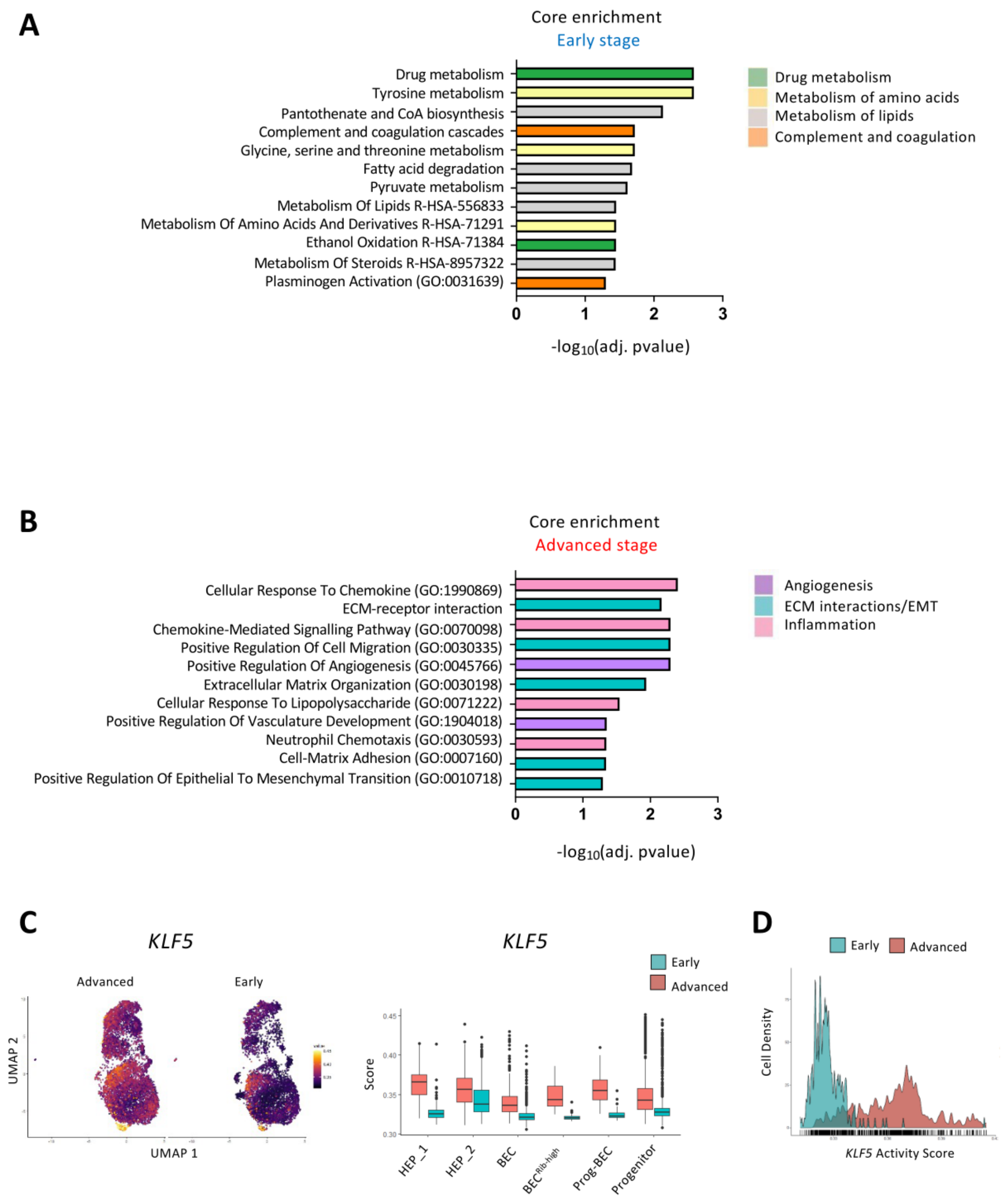


**Supplementary Figure 5. b-Orgs$^{HEP}$ recapitulate disease stage features. Related to Figure 5.**

**(A) and (B)** Significantly enriched biological processes (Reactome, Gene Ontology, Kegg) in b-Orgs$^{HEP}$ and matched biopsies from early stages **(A)** or advanced stages **(B)** using core enrichment genes from GSEA analysis. Bar colours group biological processes into major terms. **(C)** On the left, UMAP visualization of predicted *KLF5* activity in early (n=1) and advanced (n=3) b-

Org$^{HEP}$ scRNA-seq data sets. On the right, box plots showing average activity score for *KLF5* in each of the non-cycling epithelial clusters forming b-Orgs$^{HEP}$, dividing samples in early and advanced ALD condition. Boxes represent interquartile range and whiskers represent minimum and maximum. Dots represent outliers. **(D)**Predicted *KLF5* regulon activity score distribution for each cell at both early and advanced disease stages. ALD, alcohol-associated liver disease; b-Orgs, biopsy-derived organoids; GSEA, gene set enrichment analysis; scRNA-seq, single-cell RNA-sequencing.

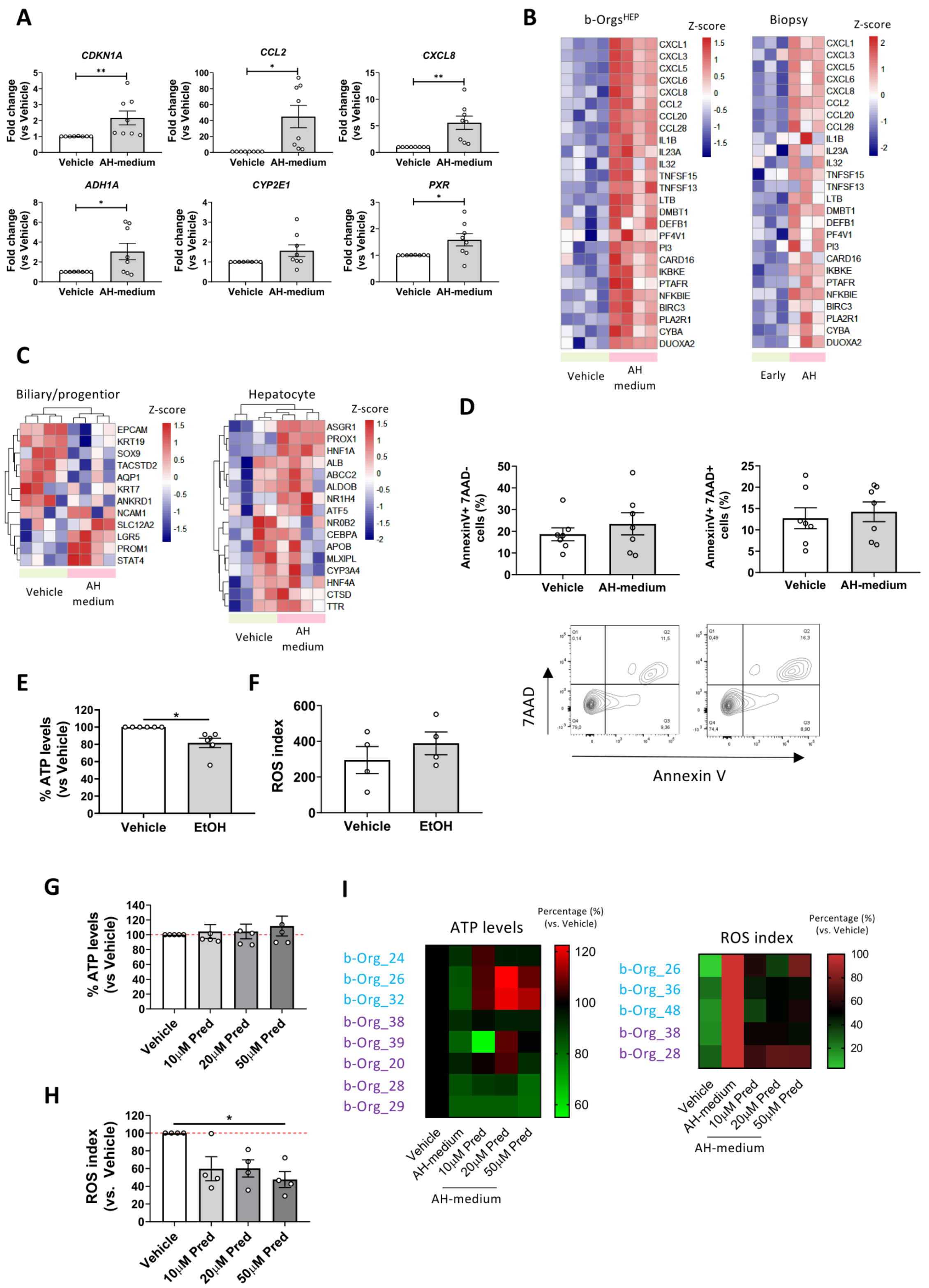


**Supplementary Figure 6. b-Orgs model ALD progression and AH pathophysiology. Related to Figure 6. (A)** Gene expression analysis of treated b-Orgs$^{HEP}$ with AH-medium compared to vehicle group (n=8 per group). **(B)**

Heatmaps showing z-score values for enriched genes involved in inflammation and oxidative stress in b-Orgs$^{HEP}$ treated with AH-medium (n=4) and AH tissue (n=3) when compared to vehicle b-Orgs (n=4) and early-stage tissue (n=3), respectively. **(C)** Heatmaps representing gene expression levels of biliary/progenitor (left) and hepatocyte (right) markers in AH-medium treated b-Orgs$^{HEP}$ (n=4) and vehicle conditions (n=4). **(D)** Annexin V and 7AAD levels detected by flow cytometry in EPCAM+ cells in vehicle and AH-medium treated b-Orgs$^{HEP}$ (n=7). **(E)** Percentage (%) of ATP levels in b-Orgs$^{HEP}$ treated with EtOH for 5 days after normalization to paired vehicle sample (n=6). **(F)** ROS index in EtOH treated and untreated b-Orgs$^{HEP}$. Quantification of ROS index was determined as ROS+ alive cells multiplied by ROS median intensity of alive cells (n=4). **(G)** Percentage (%) of ATP levels in b-Orgs$^{HEP}$ treated for 3 days with different concentrations of Prednisolone. ATP levels are normalized to vehicle group (n=4). **(H)** ROS index in b-Orgs$^{HEP}$ treated with different concentrations of Prednisolone for 3 days. Data is normalized to the vehicle group (n=4). **i.** Heatmaps depicting individual response of each b-Org$^{HEP}$ line to 5-day treatment with AH-medium and different concentrations of Prednisolone. Scale colour corresponds to percentage (%) of ATP levels normalized to vehicle group (top, n=8) or ROS index related to AH-medium (bottom, n=5). b-Org$^{HEP}$ lines generated from early stage are shown in blue, and the ones from advanced stage are shown in purple. All data is presented as mean ± SEM; no significance, * $p< 0.05$, ** $p < 0.01$, was determined by paired t-test (for *CCL2, ADH1A, CYP2E1 and PXR* in A, D and F), Wilcoxon matched-pairs signed rank test (for *CDKN1A* and *CXCL8* in A and E), and Friedman test with Dunn's multiple comparisons test (G and H).

AH, alcohol-associated hepatitis; b-Orgs, biopsy-derived organoids; EtOH, ethanol; Pred, Prednisolone; ROS, reactive oxygen species.

**Supplementary Table S1.** Baseline characteristics of patients included in the study.

| Variables | Early stages (n=28) | Advanced stages (n=34) |
|---|---|---|
| Sex, female | 7 (25) | 8 (24) |
| Age, years | 59 (50-64) | 55 (44-63) |
| Fibrosis stage, METAVIR | | |
| • F0-F1 | 12 (43) | 1 (3) |
| • F2-F3 | 11 (39) | 6 (18) |
| • F4 | 5 (18) | 27 (79) |
| AST, UI/L | 48 (26-97) | 111 (70-152) |
| GGT, UI/L | 185 (100-519) | 220 (71-692) |
| Bilirubin, mg/dL | 0.7 (0.5-1.2) | 12.4 (4.7-18.8) |
| Albumin, g/L | 42 (37-46) | 29 (27-32) |
| INR | 1.1 (1.0-1.1) | 1.7 (1.3-2.0) |
| MELD score | 7 (6-10) | 22 (17-28) |

Abbreviations: AST, aspartate aminotransferase; GGT, gamma-glutamyltransferase; INR, international normalized ratio; MELD, model for end-stage liver disease; METAVIR, meta-analysis of histological data in viral hepatitis.

**Supplementary Table S2.** Excel file containing cluster DEGs from b-Orgs$^{HEP}$ scRNA-seq. Related to Figure 4.

**Supplementary video 1. Organoids budding from a biopsy embedded in Matrigel. Related to Figure 1.** Time lapse showing how organoids emerge and grow from a biopsy. The video covers growth from day 2 after plating the biopsy to day 5. Scale bar = 0.08 cm.

**Supplementary video 2. MRP2 and E-cadherin staining in b-Orgs$^{HEP}$. Related to Figure 3.** Video showing successive Z-stacks from a b-Org$^{HEP}$ stained for MRP2 (green) and E-cadherin (red).
